# Analyzing combined eye-tracking/EEG experiments with (non)linear deconvolution models

**DOI:** 10.1101/735530

**Authors:** Olaf Dimigen, Benedikt V. Ehinger

## Abstract

Fixation-related potentials (FRPs), neural responses aligned to saccade offsets, are a promising tool to study the dynamics of attention and cognition under natural viewing conditions. In the past, four methodological problems have complicated the analysis of such combined eye-tracking/EEG experiments: (i) the synchronization of data streams, (ii) the removal of ocular artifacts, (iii) the condition-specific temporal overlap between the brain responses evoked by consecutive fixations, (iv) and the fact that numerous low-level stimulus and saccade properties also influence the post-saccadic neural responses. While effective solutions exist for the first two problems, the latter ones are only beginning to be addressed. In the current paper, we present and review a unified framework for FRP analysis that allows us to deconvolve overlapping potentials and control for linear and nonlinear confounds on the FRPs. An open software implementation is provided for all procedures. We then demonstrate the advantages of this approach for data from three commonly studied paradigms: face perception, scene viewing, and natural sentence reading. First, for a traditional ERP face recognition experiment, we show how deconvolution can separate stimulus-ERPs from overlapping muscle and brain potentials produced by small (micro)saccades on the face. Second, in scene viewing, we isolate multiple non-linear influences of saccade parameters on the FRP. Finally, for a natural sentence reading experiment using the boundary paradigm, we show how it is possible to study the neural correlates of parafoveal preview after removing spurious overlap effects caused by the associated difference in average fixation time. Our results suggest a principal way of measuring reliable fixation-related brain potentials during natural vision.

## INTRODUCTION

During everyday life, we make two to four eye movements per second to extract new information from our visual environment. Despite the fundamentally active nature of natural vision, the electrophysiological correlates of visual cognition have mostly been studied under passive viewing conditions that minimize eye movements. Specifically, in most event-related potential (ERPs) experiments, participants are instructed to fixate the screen center, while stimuli are presented at a comparatively slow pace.

An alternative approach that has gained popularity in recent years are simultaneous recordings of eye movements with the electroencephalogram (EEG) during the free viewing of complex stimuli. In such co-registration studies, the EEG signal can then be aligned to the end of naturally occurring eye movements, yielding fixation-related brain potentials (FRPs, for reviews see Baccino, 2011; Dimigen, Sommer, Hohlfeld, Jacobs, & Kliegl, 2011; Nikolaev, Meghanathan, & van Leeuwen, 2016; Velichkovsky et al., 2012). Compared to traditional passive stimulation paradigms without eye movements, this data-rich technique has the advantage that it combines the behavioral information gained from eye tracking (such as fixation durations and locations) with the high temporal resolution and neurophysiological markers provided by EEG, allowing the researcher to resolve the attentional, cognitive, and affective processes occurring within individual fixations. Due to the rapid pace of saccade generation during natural vision, it is also potentially possible to collect up to 10,000 evoked responses per hour under ecologically valid conditions.

However, the co-registration of eye movements and EEG during free viewing is also complicated by several data-analytic challenges (Dimigen et al., 2011) which have hampered the more widespread adoption of this technique in neurocognitive research. These problems, as illustrated in Figure 1, are (1) the synchronization and integration of the two data streams, (2) the ocular measurement artifacts caused by movements of the eye balls, eye lids, and extraocular muscles, (3) the condition-specific temporal overlap between the brain responses evoked by successive fixations, and (4) the strong and often nonlinear influences of visual and oculomotor low-level variables on the neural responses produced by each eye movement. Whereas there are decent solutions for the first two problems, the latter two are only beginning to be solved. In the current paper, we describe an integrated analysis framework for EEG analyses during natural vision which addresses these remaining two problems (overlap and low-level influences). We also provide a tutorial review on how this framework can be implemented using a recently introduced open-source toolbox that offers all of the necessary procedures (Ehinger & Dimigen, 2019). Finally, to demonstrate how this approach can improve the analysis of unconstrained viewing experiments and produce new theoretical insights, we will apply it to co-registered datasets from three domains of neurocognitive research: face perception, scene viewing, and sentence reading.

**Figure 1.**
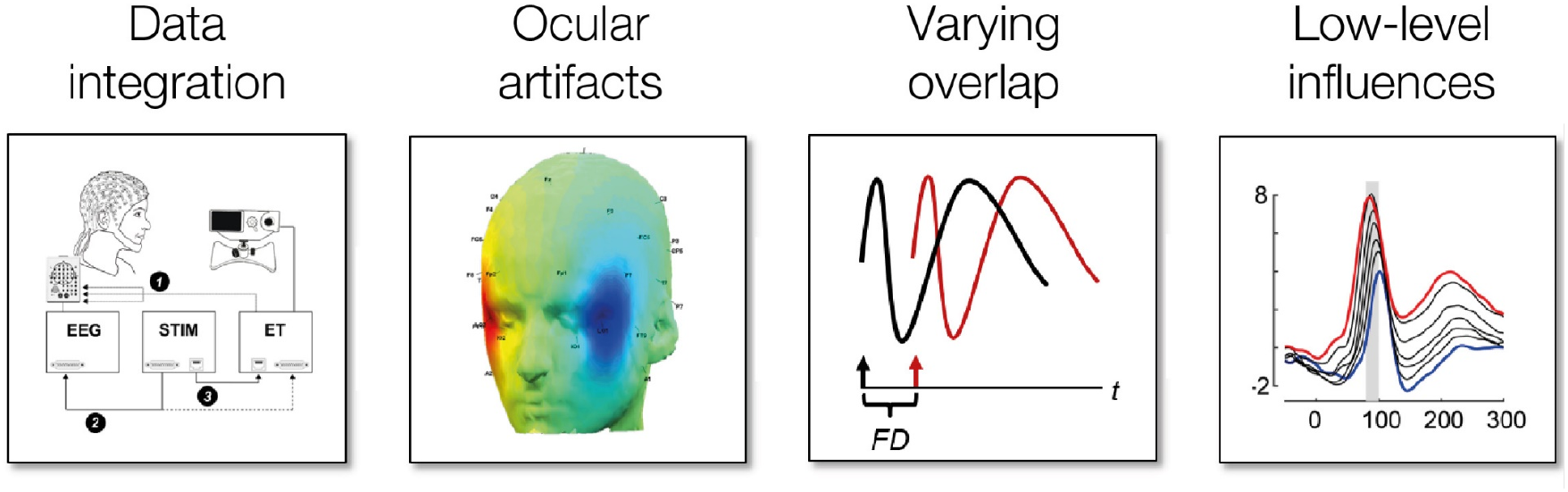
Schematic overview of four data-analytic challenges encountered when co-recording eye movement and EEG data during natural vision (after Dimigen et al., 2011). Whereas good solutions already exist for the first two problems, deconvolution modeling with nonlinear predictors effectively addresses the problems of varying overlap and low-level influences on the brain signals.

### Four problems related to free viewing

The four methodological problems are illustrated in Figure 1: The first problem summarizes several technical issues related to the simultaneous recording, precise synchronization, and joint representation of the EEG and the video-based eye-tracking data. Nowadays these issues are largely solved by optimizing the laboratory setup and by sending shared trigger pulses to both systems during each trial (e.g. Baccino & Manunta, 2005). The two recordings can then be aligned offline at millisecond precision with existing software (e.g. the EYE-EEG toolbox; Dimigen et al., 2011; see also Baekgaard et al., 2014; Xue et al., 2017) that also adds saccade and fixation onsets as additional event markers to the EEG.

A second and more critical problem are the strong EEG artifacts generated by movements of the eye balls, eyelids, and extraocular muscles during free viewing (Keren, Yuval-Greenberg, & Deouell, 2010; Lins, Picton, Berg, & Scherg, 1993; Plöchl, Ossandón, & König, 2012). The eye balls, in particular, act as electrostatic dipoles that rotate with each eye movement producing large voltage distortions across the scalp (corneoretinal artifact). Two smaller artifacts are produced by the relative movement of the eye lids over the cornea during upward saccades (Lins et al., 1993; Plöchl et al., 2012) and by the recruitment of the eye muscles at saccade onset (saccadic spike potential, Blinn, 1955; Keren et al., 2010; Yamazaki, 1968). All three ocular artifacts – corneoretinal, eye lid, and spike potential – need to be removed from the EEG without distorting brain activity. Algorithms like independent component analysis (ICA, Jung et al., 1998) have commonly been used to suppress ocular artifacts, even under free viewing conditions (Henderson, Luke, Schmidt, & Richards, 2013; Hutzler et al., 2007; Keren et al., 2010; Leeuwen & Nikolaev, 2018; Plöchl et al., 2012; Ries, Slayback, & Touryan, 2018b). Although some residual artifacts (in particular from spike potentials) are visible in the waveforms of most published FRP studies, it seems likely that correction procedures can be improved further in the future, for example, by taking into account the information provided by the eye-tracker. Specifically, the concurrent eye-tracking data is useful to select optimal training data for the ICA algorithm (Craddock, Martinovic, & Müller, 2016; Keren et al., 2010), to identify artifact components (Plöchl et al., 2012), and to evaluate the results of the correction (Dimigen, 2020; Ries et al., 2018b). Recent findings indicate that if ICA procedures are fine-tuned for free viewing experiments in this manner, ocular artifacts can be almost fully removed with relatively little distortion of genuine brain activity (Dimigen, 2020; Ries et al., 2018b).

The last two major problems, differences in temporal overlap and low-level covariates, have received less attention but are a direct consequence of the fast pace and quasi-experimental nature of normal visual exploration behavior. In traditional EEG laboratory experiments, the experimenter has full control over the timing and sequence of the presented stimuli and the participant’s motor behavior is often restricted to a single button press. In most cases, it is also possible to match the visual low-level properties of the stimuli between conditions. In contrast, in any experiment with multiple saccades, the participant rather than the experimenter decides where to look and when to look at a stimulus belonging to a given condition. This means that the durations of fixations, the size and direction of saccades, and the low-level features of the stimulus at the currently foveated location (e.g. the local luminance and local contrast of an image) are not only intercorrelated with each other (Nuthmann, 2017), but usually different between the experimental conditions. Since all of these factors also influence the post-saccadic brain response, this can easily lead to condition-specific distortions and incorrect conclusions. In the following sections, we will describe these two problems and some proposed solutions in more detail.

### Varying temporal overlap

Event-related potentials that index cognitive processes often last up to a second before the signal tapers off and returns to baseline. In contrast, the average fixation lasts only about 200-400 ms (Rayner, 2009) in most viewing tasks. This rapid pace of natural vision means that brain responses elicited by any given fixation will strongly overlap with those of preceding and following fixations. While this overlapping activity is smeared out due to the normal variation in fixation durations, overlap becomes a problem whenever the distribution of fixation durations differs between conditions.

As an example, consider a visual search task, where participants look for a target item within a complex visual scene while their EEG is recorded (Cooper et al., 1977). It is well-known that on average, task-relevant target items are fixated longer than irrelevant non-targets (e.g. Brouwer, Reuderink, Vincent, van Gerven, & van Erp, 2013). This in turn means that the visually-evoked lambda response (the analogue of the P1 component in FRPs) from the next fixation will overlap at an earlier latency in the non-target condition than in the target condition (see also Figure 2C for a similar example). Unless special precautions are taken (e.g. as in Kamienkowski, Ison, Quiroga, & Sigman, 2012), cognitive effects of target processing on the P300 component will therefore be confounded with trivial condition differences produced by the change in overlap. In other words, any difference in mean fixation duration between conditions can produce spurious differences in the EEG, even if the real underlying brain activity is the same in both conditions. Furthermore, if the duration of the pre-target fixation differs between conditions (e.g. because of the extrafoveal preprocessing of the target), these distortions will also affect early parts of the FRP waveform, including the baseline interval before fixation onset (see also Figure 7D below). The confounding effect of overlapping potentials is also illustrated in Figure 2 for a simulated experiment with cars and faces.

**Figure 2.**
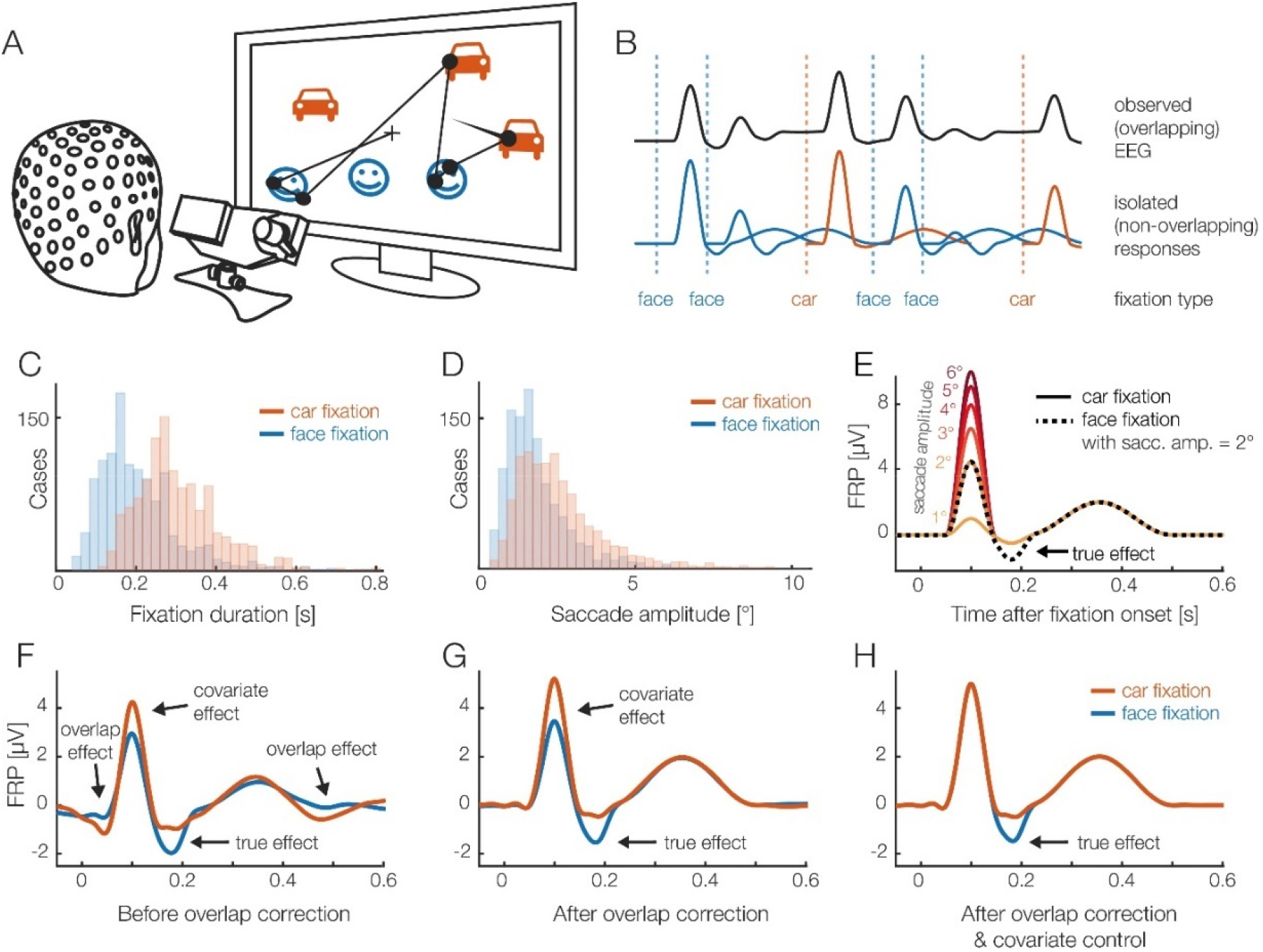
(A) A hypothetical eye-tracking/EEG experiment (simulated data) in which participants freely looked at pictures of faces and cars. We are interested whether the fixation-related potential (FRP) differs between stimulus categories. (B) Short interval of the recorded EEG. Every fixation on a car or face elicits a brain response (lower row). Because brain responses last longer than the respective fixation durations, we can only observe the sum of the overlapping responses in the EEG (upper row). (C) In most free viewing tasks, eye movement behavior differs systematically between conditions. For this example, we assume that on average, faces are fixated more briefly than cars. This means that the overlap with the brain responses of the next fixations is stronger in the face condition. (D) Furthermore, we assume that due to more refixations on faces, saccades are on average smaller in the face condition. (E) We also assume that there is a true effect: the N170 component of the FRP is larger for faces than cars (Rossion & Jacques, 2012). In addition, however, it is well-known that saccade size alone has a strong nonlinear effect on the lambda response, the P1 component of the FRP. (F) Average FRP that would be measured in each condition. In addition to the genuine N170 effect, this simulation reveals several spurious effects, caused by the differences in fixation duration and saccade size. (G) Linear deconvolution corrects for the effects of overlapping potentials. (H) To also remove the confounding effect of saccade amplitude, we need to also include saccade amplitude as a (nonlinear) predictor in the model. (H) Now we can recover just the true N170 effect free of confounds. A conceptually similar figure was used in Ehinger & Dimigen (2019).

A second, but frequently overlooked type of overlap is that between stimulus-onset ERPs and FRPs. In most free viewing experiments, a single visual stimulus – for example a search array, a sentence, or a scene – is presented at the beginning of each trial. The fixation-related potentials therefore not only overlap with each other, but also with this stimulus-ERP, which is often strong and long-lasting (cf. Figure 1E in Dimigen et al., 2011). This means that fixations that happen early and late during a trial differ systematically in terms of their baseline activity and cannot be directly compared to each other. In practice, this type of overlap can be just as problematic as that from neighboring fixations (Coco, Nuthmann, & Dimigen, 2020; de Lissa et al., 2019).

Finally, overlapping potentials are also relevant in traditional EEG experiments in which the stimulus-ERP is the signal of interest, and eye movements are just a confound. In particular, potentials from involuntary (micro)saccades executed during the trial have been shown to distort stimulus-locked EEG analyses in the frequency (Yuval-Greenberg, Tomer, Keren, Nelken, & Deouell, 2008) and time domain (Dimigen, Valsecchi, Sommer, & Kliegl, 2009). We therefore need effective methods to disentangle overlapping activity from multiple events. Several workarounds to the overlap problem have been proposed that center on data selection. One simple approach to reduce overlap is to analyze only long fixations with a minimum duration (e.g. > 500 ms) (e.g. Brouwer et al., 2013; Kamienkowski, Ison, Quiroga, & Sigman, 2012; Kaunitz et al., 2014); another is to analyze only the first or last fixation in a sequence of fixations, which eliminates overlapping activity from the preceding and subsequent fixation, respectively (Hutzler et al., 2007). Of course, solutions like these are not optimal because they either exclude a large portion of the data or place strong constraints on the possible experimental designs.

A stark improvement is found in the deconvolution approach that was first used for EEG analyses in the 1980s (Eysholdt & Schreiner, 1982; Hansen, 1983). Here, the measured continuous EEG signal is understood as the convolution of the experiment events (i.e., a vector which contains impulses at the latencies of the experimental events) with the isolated brain responses generated by each type of event (as illustrated in Figure 2B). The inverse operation is deconvolution, which recovers the unknown isolated brain responses given only the measured (convolved) EEG signal and the latencies of the experimental events (Figure 2G; Ehinger & Dimigen, 2019). Deconvolution is only possible because the events show different temporal overlap with each other; conversely, without any jitter (i.e., with a constant stimulus-onset asynchrony or SOA), the responses would be inseparable because it is ambiguous whether the observed activity was evoked by the current event or the preceding event. This can be achieved by experimentally adding jitter to event onset times, or it happens naturally as fixation durations vary between fixations. Such temporal variability allows us to recover the unknown isolated responses, under two assumptions: (1) the brain signals evoked by different events add up linearly and (2) the degree of temporal overlap between the events does not change the processing in the brain itself and the neural response evoked by each event. The first assumption is met due to the linear summation of electrical fields (Nunez & Srinivasan, 2006). The second assumption – that the underlying brain responses are the same, regardless of the amount of overlap in the experiment – is likely incorrect, but in practice still a useful approximation (see *Discussion*).

Early iterative deconvolution techniques for EEG (in particular the ADJAR algorithm, Woldorff, 1993) have proven difficult to converge (Kristensen, Rivet, & Guérin-Dugué, 2017; Talsma & Woldorff, 2004), do not allow to simultaneously control for the influences of continuous covariates on the EEG, or were designed for specialized applications (Ouyang et al., 2011). More recently, a linear deconvolution method based on the least-squares estimation was successfully applied to solve the overlap problems in EEG (Burns, Bigdely-Shamlo, Smith, Kreutz-Delgado, & Makeig, 2013; Cornelissen, Sassenhagen, & Võ, 2019; Dandekar, Privitera, Carney, & Klein, 2012; Guérin-Dugué et al., 2018; Kristensen, Guerin-Dugué, & Rivet, 2017; Kristensen, Rivet, et al., 2017; Litvak, Jha, Flandin, & Friston, 2013; Lütkenhöner, 2010; Sassenhagen, 2018; N. J. Smith & Kutas, 2015b; Spitzer, Blankenburg, & Summerfield, 2016). This approach, initially applied to fMRI (Dale & Buckner, 1997; Glover, 1999; Serences, 2004), has crucial advantages over the previous, often iterative approaches. The properties of the linear model are not only well understood, but the embedding of deconvolution in the linear model allows for multiple regression, meaning that many different event types (like stimulus onsets, fixation onsets, and button presses) can be modeled within the same model together with continuous covariates. We provide a non-mathematical review of this linear deconvolution approach in the *Tutorial Review* section further below.

### Low-level covariates influencing eye movement-related responses

After adequate correction for overlap, only the fourth, serious problem remains: the massive influence of visual and oculomotor low-level variables on the shape of the saccade- or fixation-related brain responses. As an example, consider the lambda response, the dominant visually-evoked component of the FRP that peaks at occipital scalp sites at around 80-100 ms after fixation onset. The lambda response is the equivalent of the P1 in FRPs, and like the P1, it is generated in striate and/or extrastriate visual cortex (Dimigen et al., 2009; Kazai & Yagi, 2003). Like much of the rest of the FRP waveform (see also Figure 7E further below), the lambda response is strongly influenced by the size of the incoming saccade (Armington & Bloom, 1974; Dandekar, Privitera, et al., 2012; Dimigen et al., 2011; Kaunitz et al., 2014; Ries et al., 2018b; Thickbroom, Knezevič, Carroll, & Mastaglia, 1991). If saccade amplitudes differ between conditions, an analogous problem to the previously discussed fixation duration bias will occur – the experimental condition with larger saccades will also have larger lambda responses. Increasing saccade amplitude also increases the amplitude of the pre-saccadic cortical motor potentials that ramp up slowly before eye movement onset in saccade-onset locked ERPs (Becker, Hoehne, Iwase, & Kornhuber, 1972; Everling, Krappmann, & Flohr, 1997; Herdman & Ryan, 2007; Richards, 2003). Due to these premotor potentials, saccade amplitude can also affect the typical pre-fixation baseline interval for the FRP (Nikolaev, Jurica, Nakatani, Plomp, & Leeuwen, 2013; Nikolaev et al., 2016).

Other visual and oculomotor covariates will introduce similar biases. For example, both the stimulus features in the currently foveated image region (Dimigen, Sommer, & Kliegl, 2013; Gaarder, Krauskopf, Graf, Kropfl, & Armington, 1964; Kristensen, Rivet, et al., 2017; Ossandon, Helo, Montefusco-Siegmund, & Maldonado, 2010; Ries et al., 2018b), the fixation location on the screen (Cornelissen et al., 2019; Dimigen et al., 2013), and the direction of the incoming saccade (Cornelissen et al., 2019; Meyberg, Werkle-Bergner, Sommer, & Dimigen, 2015; see also the results in this paper) modulate the FRP waveshape. It is therefore reasonable to conclude that most existing FRP results (including our own) are confounded to some degree, since they did not fully control for differences in overlap and low-level covariates.

### Fixation matching: Limitations and problems

One proposed method to partially control for these confounding factors is post-hoc fixation matching (Dias, Sajda, Dmochowski, & Parra, 2013; Kamienkowski et al., 2012; Leeuwen & Nikolaev, 2018; Luo, Parra, & Sajda, 2009). The underlying idea is simple: After the experiment, the researcher selects those fixations from each experimental condition that are the most similar in terms of overlap and a few of the most important visuomotor covariates (e.g. incoming saccade amplitude) by selecting them based on their distance to each other in this multi-dimensional feature space (e.g. the Mahalanobis distance in Nikolaev et al., 2016). These matched subsets of fixations are then compared, while the remaining fixations are discarded. After matching, the oculomotor covariates are as similar as possible across conditions and all conditions are affected by overlap to a similar degree. To describe it differently, with fixation matching, we attempt to convert a quasi-experimental, naturalistic situation (free viewing) back into an orthogonal, well-controlled experiment.

Matching procedures are relatively easy to implement and if the durations of the fixations in the different conditions are also matched, they also address the overlap problem. However, there are also several limitations to this method. First, there is a loss of data due to fixations that cannot be matched. Second, the number of covariates that can be simultaneously matched under practical conditions is limited and likely smaller than the number of variables that are already known to modulate the FRP. In particular, variables shown to affect the waveform include the durations of the preceding and current fixation, the temporal overlap with the stimulus-onset ERP (Coco et al., 2020), the amplitude (e.g., Gaarder et al., 1964; Thickbroom et al., 1991) and direction (Cornelissen et al., 2019; Meyberg et al., 2015) of the incoming saccade, the fixated region of the screen (Dimigen et al., 2013), and local image properties at the foveated spot, such as the local luminance (Armington, Gaarder, & Schick, 1967; Gaarder et al., 1964; Kristensen, Rivet, et al., 2017; Ossandon et al., 2010) and spatial frequency spectrum (Armington & Bloom, 1974; Armington et al., 1967; Ries et al., 2018b; Yagi, Ishida, & Katayama, 1992). In our experience, in a task like sentence reading, it is rarely possible to match more than two to three covariates, at least if these covariates show sizeable condition differences in the first place. Third, matching approaches are limited to simple factorial designs; it is hard to imagine how a matching procedure would work if the independent variable manipulated in the experiment (e.g. the saliency of an image region) would be continuous rather than categorical in nature. Fourth, currently proposed algorithms for fixation matching are based on predefined thresholds or null-hypothesis testing. In the latter case, it is assumed that there is no longer a difference in the mean of the covariates if the difference is not statistically significant anymore. However, a non-significant difference between the covariates after matching does not mean that the null hypothesis is correct (Sassenhagen & Alday, 2016).

Finally, it is possible that the matching of saccade or fixation properties reduces the actual psychological effect in the data. For example, once fixation durations are matched, we are comparing the neural correlates of two pools of fixations that on average did *not* differ in terms of the behavioral outcome. In contrast, the fixations that may be the psychologically most relevant ones that contribute the most to the behavioral effect – those at the tail of the distribution – tend to be eliminated from the FRP by the matching. Discarding these fixations could therefore lead to more false negative findings.

### Towards a unified model based on linear deconvolution and spline-regression

Based on these problems, it is clear that another solution needs to be found. Instead of selecting fixations, we need to correct for the aforementioned effects. One tool to account for these confounds is multiple linear regression with continuous regressors. The approach, called mass-univariate linear modeling (Figure 3), has been frequently applied to traditional EEG datasets (Amsel, 2011; Hauk, Davis, Ford, Pulvermüller, & Marslen-Wilson, 2006; Pernet, Chauveau, Gaspar, & Rousselet, 2011; Rousselet, Pernet, Bennett, & Sekuler, 2008) and more recently also to account for linear influences of saccade parameters on FRPs (Weiss, Knakker, & Vidnyánszky, 2016). Importantly, the linear modeling of covariates has recently also been shown to combine seamlessly with the linear deconvolution approach introduced above, both for ERPs (N. J. Smith & Kutas, 2015b) and FRPs (Cornelissen et al., 2019; Ehinger & Dimigen, 2019; Guérin-Dugué et al., 2018; Kristensen, Rivet, et al., 2017).

**Figure 3.**
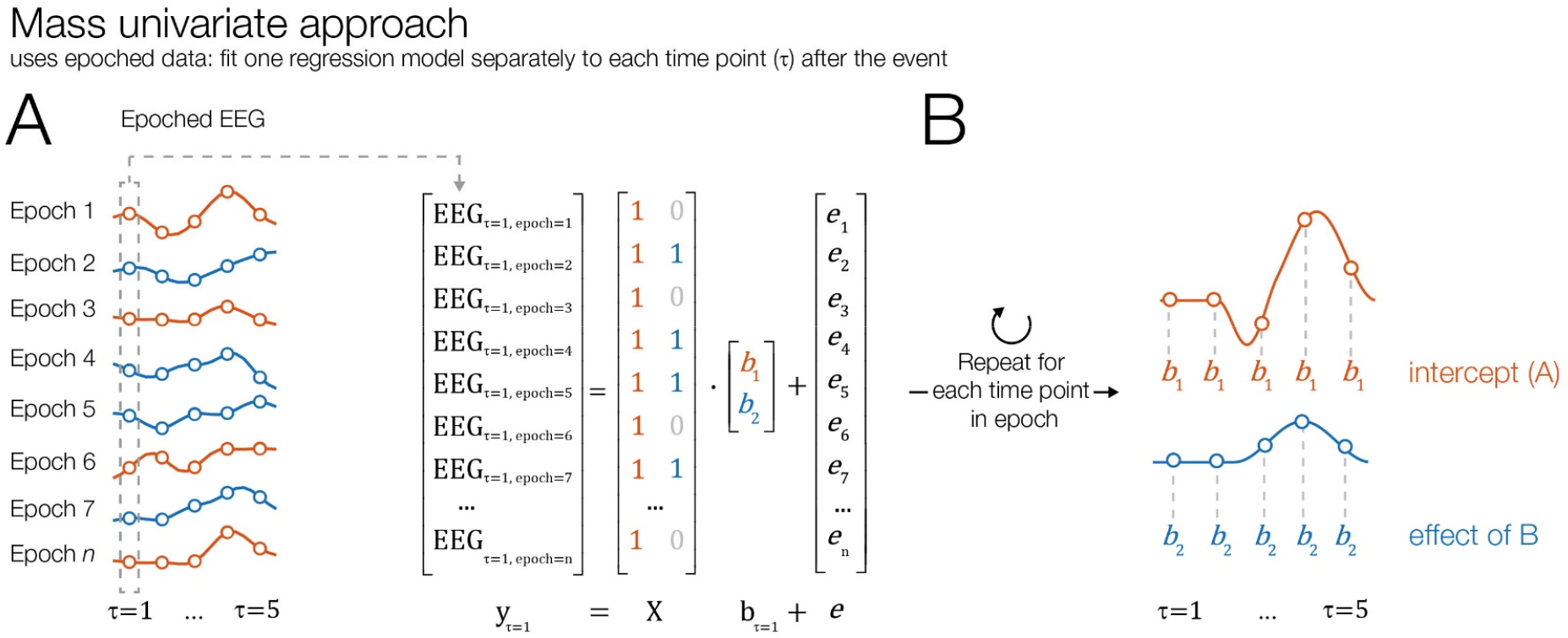
Illustration of mass univariate modeling of EEG (without overlap correction). (A) Similar to a traditional ERP analysis, short epochs are cut around all events of interest (e.g., fixation onsets on stimuli of type “A” and “B”). A regression model is then fit separately to each time point within the epoch, with the EEG signal over trials as the dependent variable. The design matrix *X* codes the value of the predictors in each trial. In this example, we model each time point by an intercept term, which captures the overall brain response generated by events of type “A” (assuming treatment coding). A second predictor only captures the additional effect elicited by events of type “B”. (B) After the model is repeatedly run for each time point, the resulting regression coefficients (*betas*) can be plotted as an ERP-like waveform (*rERP*). The mass univariate approach can also account for covariates, but not for overlapping potentials from preceding and following events.

A fact that complicates the problem even further is that the relationship between saccade properties and the FRP is often nonlinear. For example, several studies found that with increasing saccade amplitude, the lambda response increases in a nonlinear fashion (Dandekar, Privitera, et al., 2012; Kaunitz et al., 2014; Ries et al., 2018b; Thickbroom et al., 1991; Yagi, 1979). Nonlinear relationships with saccade size have also been reported for the saccadic spike potential (Armington, 1978; Boylan & Doig, 1989; but see also Keren et al., 2010), the burst of eye muscle activity at saccade onset. As we will confirm below, the influences of some oculomotor covariates on the FRP are indeed highly nonlinear. Ignoring these nonlinear relations and modeling the data by a simple linear predictor can therefore produce suboptimal results and bias the results in an arbitrarily way (e.g. Tremblay and Newman, 2015).

Fortunately, due to the flexible nature of the linear model, it is also possible to model nonlinear relationship*s* within this framework. For this purpose, the linear model is augmented by spline regression, as used in the generalized additive model (GAM, Wood, 2017). Recently, spline regression has been applied to ERPs (Hendrix, Baayen, & Bolger, 2017; Kryuchkova, Tucker, Wurm, & Baayen, 2012; Tremblay, A., & Baayen, 2010; Tremblay & Newman, 2015) and also to FRPs (Van Humbeeck, Meghanathan, Wagemans, van Leeuwen, & Nikolaev, 2018). In this paper, we demonstrate how spline regression can be combined with deconvolution to control for the highly nonlinear influences of some predictors (exemplified here for saccade amplitude and saccade direction) during the free viewing of pictures.

### Current paper

Combining the ideas presented above, we propose that the combination of linear deconvolution with nonlinear spline regression (GAM) can solve both of the major remaining problems, overlap and confound control. In the remainder of this paper, we will first describe both methods on an intuitive level by building up a model for a typical free viewing experiment, step by step. To illustrate the advantages of this approach on real data, we then use the recently introduced *unfold* toolbox (http://www.unfoldtoolbox.org; Ehinger & Dimigen, 2019) to analyze combined eye-tracking/EEG data from three paradigms: face recognition, visual search in scenes, and reading.

## (NON)LINEAR DECONVOLUTION: A TUTORIAL REVIEW

In this section, we first review the basic principles of deconvolution modeling within the linear regression framework. Afterwards, we will outline how deconvolution combines with the concept of spline predictors to model nonlinear effects. Introductions to linear models, regression-based ERPs (rERP), and deconvolution within the rERP framework can be found in Smith and Kutas (2015a, 2015b), Ehinger & Dimigen (2019), and Sassenhagen (2018). Recent applications to saccade- and fixation-related potentials are found in Coco et al. (2020), Cornelissen et al. (2019), Dandekar et al. (2012), Guérin-Dugué et al. (2018) and Kristensen et al. (2017b). A more technical description of some of the steps summarized below is provided in Ehinger & Dimigen (2019). In the following, we restrict ourselves to a description of the general principles in a less technical and more intuitive fashion.

### Linear modeling of the EEG

Before we introduce linear deconvolution, let us first look at another common way that multiple regression is applied to EEG: Mass-univariate linear modeling (Pernet et al., 2011). Figure 3A illustrates this approach for a simple experiment with two conditions, but the concept can be generalized to arbitrary designs. In a first step, we cut the data into epochs around the onsets of experimental events, for example fixations (e.g. Weiss et al., 2016). For each of the *n* time points in the epoch, we then fit a separate regression model, which tries to explain the observed data over trials at the given time point *t*:

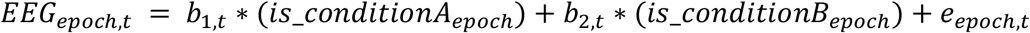

The same model can also be written in matrix notation:

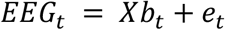

Here, *X* is the design matrix. Each of its rows represents one observation (e.g. one fixation onset on a target object) and each of its columns represents one predictor and its value in the respective trial (e.g., the type of object fixated or the incoming saccade size). *b* is a vector of to-be-estimated parameters (regression coefficients or *betas*) that we wish to estimate and *e* is the error term, a vector of residuals. For each time point relative to the event, we run the model again and estimate the respective betas. The result is a series of betas for each predictor, also called a regression-ERP (rERP), that can be plotted against time just like a traditional ERP waveform. Mass-univariate models has been successfully applied to ERP studies in which many covariates affect the neural response (Amsel, 2011; Hauk et al., 2006; Rousselet et al., 2008), but they cannot account for varying temporal overlap between neighboring events. In other words, if overlap differs between conditions, they will produce biased estimates.

### Deconvolution within the linear model

The linear deconvolution approach – as illustrated in Figure 4 – directly addresses this issue of overlapping brain responses. As an example, consider the left part of Figures 4A, which show a small part of a continuous EEG recording. We can see that the EEG recorded at sample 25 of the experiment (grey dashed box) is the sum of the responses to three different experimental events: the early part of the brain response to an event of type A which also happened at this sample (highlighted in orange),, the late part of the response to an event of type B (which had occurred two samples earlier, in blue) and the late part of the response to a second event of type A (which had occurred four samples earlier, in orange). During free viewing, the temporal overlap with neighboring fixation events is slightly different for each fixation. Additionally, in many experiments, participants look at different categories of stimuli during each trial, leading to variable sequences of events. For example, in the hypothetical car/face experiment depicted in Figure 2, fixations on cars are sometimes followed by a fixation on a face and sometimes by a fixation on another car. Due to both sources of variability – in terms of temporal overlap, the sequence of events, or both – it is possible to recover the nonoverlapped signals in a regression model.

**Figure 4.**
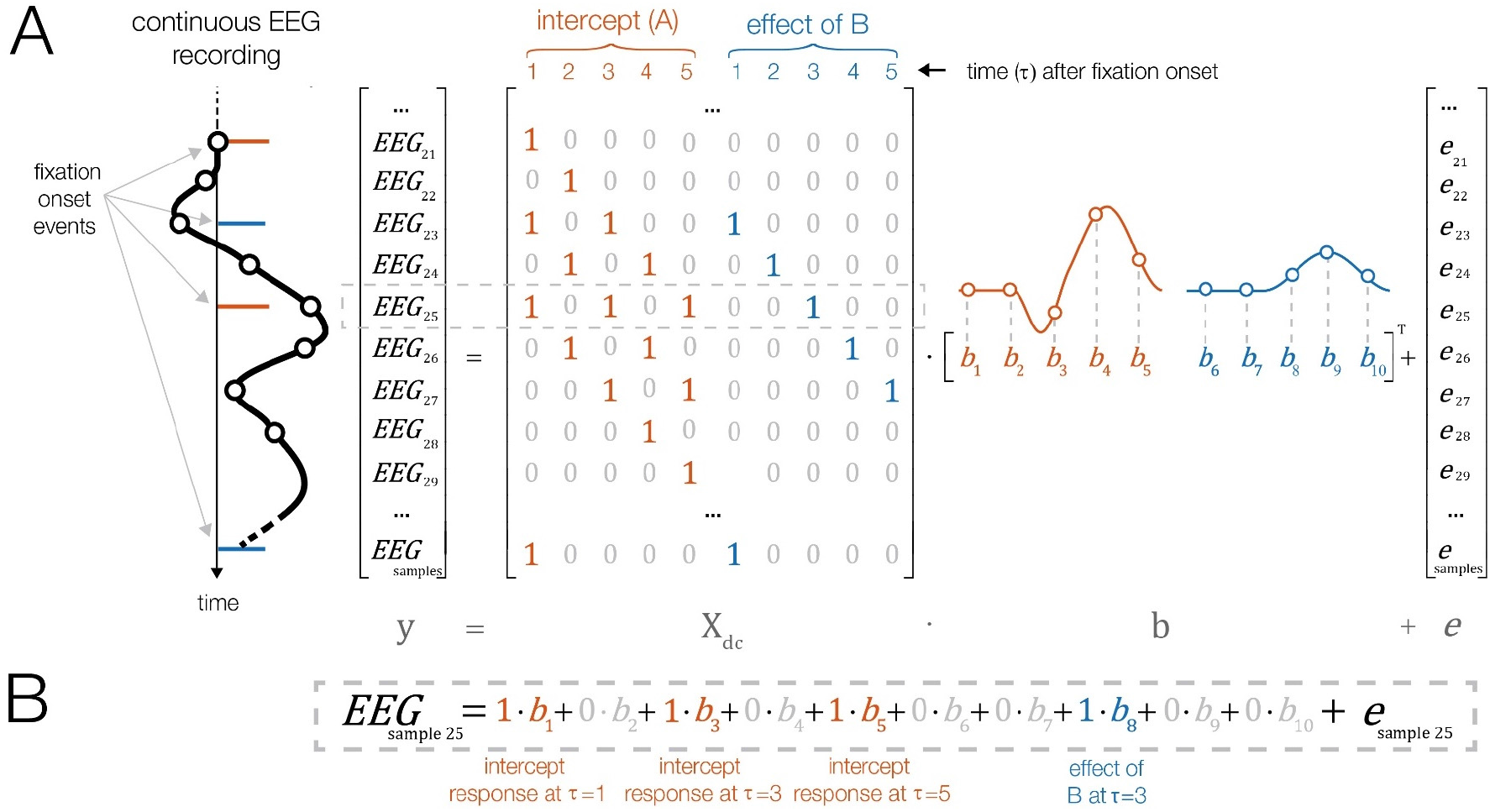
Illustration of linear deconvolution in a situation with overlapping brain responses. In contrast to the mass univariate approach, linear deconvolution explains the continuous EEG signal within a single regression model for each channel. (A) A short interval of the continuous EEG is plotted on the left (time runs from top to bottom). Highlighted are the onset latencies of four experimental events, belonging to condition *A* (in orange) and *B* (in blue). Our goal is to recover the unknown isolated brain responses (betas) elicited by each type of event so that together, they best explain the continuous EEG. For this purpose, we create a time-expanded version of the design matrix (X_dc_) in which a number of local time points around each event onset (here only for 5 time points) are added as predictors. We then solve this large regression model for the betas (plotted on the right). In this example, responses are modeled by an intercept term, which describes the overall neural response to events of type A. Furthermore, we model the additional effect of events of type B. (B) For example, the 25th sample of the continuous EEG can be explained by the sum of the overlapping brain responses to three events: the response to the first event of type *A* (at time point 5 after that event), the response to an event of type *B* (at time point 3 after that event) and by a another occurrence of an event of type “A” (at time point 1 after that event). Because the temporal distance between events varies throughout the experiment, it is possible to find a unique solution for the betas that best explains the measured EEG.

One property that distinguishes linear deconvolution from the mass-univariate approach is that the input EEG data needs to be continuous rather than cut into epochs. This is because for a correct estimation, we need to consider the temporal relationship between all event-related responses that happened close to each other in time. If we would cut the EEG into short epochs, the epochs would likely not contain all of the preceding and following events that also influenced the signal. If we would instead cut the EEG into very long epochs, the epochs would start overlapping with each other, meaning that some data points would enter the model multiple times, biasing the estimation. For these reasons, we need to model the continuous EEG (after it was corrected for ocular artifacts and eye movement markers were added).

We set up the deconvolution model by generating a new design matrix *X_dc_* (where *dc* stands for deconvolution), which spans all samples of the continuous EEG recording (Figure 4A). Like in the mass univariate model (Figure 3), the columns of this design matrix code the condition of the events in our model (e.g. “Is this an event of type A?”). In order to explain linear deconvolution, we need to introduce the concept of *local time* (τ), which describes the time (in samples) relative to the onset of a given type of event. In the simplified example of Figure 4, we model just the *n* = 5 time points following each event (from τ = 1 to τ = *5* after event onset). In a realistic scenario, one would model a couple of hundred sampling points before and after the fixation onset (e.g. in a time window from −200 to 800 ms). The time range should be chosen so that it captures the entire fixation-related EEG response, including oculomotor potentials that reflect saccade planning and precede fixation onset (Becker et al., 1972; Everling et al., 1997; Herdman & Ryan, 2007; Richards, 2003). To set up the deconvolution model, we then have to add *n* = 5 new predictors to the model per event. These predictors will directly model the time course of the event-related response; a process that we will call *time-expansion* and explain in the following:

The first *n* columns in the new design matrix *X_dc_* belong to the predictor A. The first column codes the first time point after that type of event occurred (τ = 1). This column will be 0 at every sample, except at those latencies of the EEG when an event of that type occurred; there, we set it to 1. The second column codes the second time point after the event, τ = 2. This column will get a 1 at all continuous EEG samples that were recorded one sample after an event of type A and zero everywhere else. We repeat this process for all five sampling points relative to each event A. Afterwards, we repeat the same for all occurrences of event B, adding five more columns to the design matrix.

If we look at the expanded design matrix produced by this process (Figure 4A), we see diagonal staircase-like patterns. We can now also immediately see which parts of the individual evoked responses contribute to the observed continuous EEG signal, as highlighted here for sample number 25. Note that the resulting time-expanded design matrix *X_dc_* is large; it has as many rows as there are samples in the EEG. The number of its columns is given by the number of predictors (here: 2) multiplied by the number of modeled time points (here: *n* = 5). Thus, to solve the regression model, we need to solve a large linear equation system. In realistic applications, the design matrix *X_dc_* often spans a few million rows (continuous EEG samples) and several tens of thousands of columns (predictors). Fortunately, this computationally difficult problem can nowadays be solved efficiently with modern algorithms for solving sparse matrices (time-expansion produces a very sparse matrix, i.e. a matrix mostly filled with zeros, as illustrated in Figure 4A). In summary, the regression formula changes little from the mass univariate model to the deconvolution model:

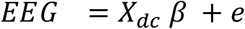

Solving this formula for *b* results in *n* betas for each event type/predictor, one for every time point modeled (see Figure 4A, right side). However, in contrast to the mass univariate model, we do not need to calculate a separate model for each time point; instead, all betas for a given EEG channel are returned in just one model fit. This time series of betas, or rERP, represent the non-overlapping brain responses for each predictor. Like traditional ERP waveforms, rERPs can be visually inspected, plotted as waveforms or scalp topographies, entered into a dipole analysis, and compared statistically (N. J. Smith & Kutas, 2015a, 2015b). Furthermore, because the estimation is linear, linear operations like a baseline correction can also be applied to the rERPs after the deconvolution (N. J. Smith & Kutas, 2015b).

### Modeling nonlinear effects

As explained above, some predictors have nonlinear influences on the EEG. As we will show in the empirical part of this paper, considering nonlinear effects is especially important in free viewing experiments. In the linear regression framework, nonlinear effects can be accounted for in different ways. Common approaches are to transform individual predictors (e.g. via a log transform) or to include higher-order terms, such as quadratic or cubic terms, in the model (polynomial regression). However, these approaches have drawbacks. The transformation of individual predictors (such as saccade amplitude) necessitates a-priori knowledge about the correct shape of the relationship, but for ERPs/FRPs, this shape is often unknown. For more complex relationships, e.g. for circular predictors with non-sinusoidal shapes, it can be difficult to find a good transformation function without resorting to an inefficient Fourier set. By using a polynomial regression one could in principle fit any arbitrary relationship, but in practice one often observes oscillatory patterns (Runge, 1901). These patterns occur because each additional term added to the polynomial acts on the entire range of the predictor, that is, it affects the fit at all saccade amplitudes, rather than just locally.

A more flexible option is spline regression, a technique also commonly summarized under the name GAM (Wood, 2017). Splines can be understood as local smoothing functions (Figure 5) and they have the advantage that they are defined locally, that is over only a short range of the continuous predictor (e.g., just for saccade amplitudes between 2° and 4°). This solves the problem of oscillatory patterns and makes the predictors easily interpretable and less dependent on the exact fit of the other parameters. The following section attempts a brief and intuitive overview of spline regression.

**Figure 5.**
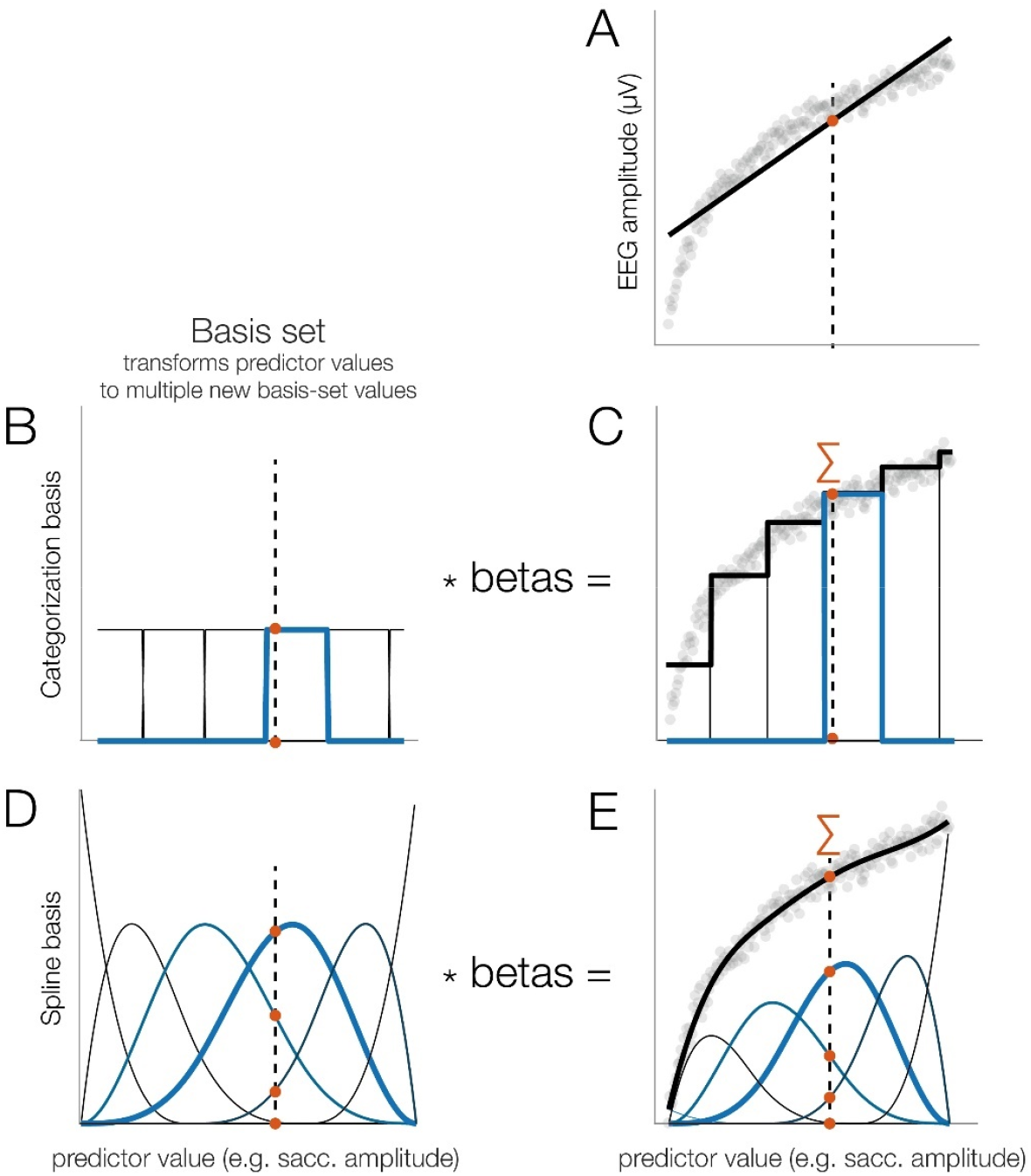
Using splines to model nonlinear effects, illustrated here for simulated data. (A) Example of a relationship between a predictor (e.g. saccade amplitude) and a dependent variable (e.g. fixation-related P1). As can be seen, a linear function (black line) fits the data poorly. The dashed vertical line indicates some example value of the independent variable (IV), e.g. a saccade amplitude of 3.1°. (B) Categorization basis set. Here, the range of the IV is “split up” into non-overlapping subranges, each coded by a different predictor which is coded as 1 if the IV for the current instance is inside the range, and as 0 otherwise. The IV is evaluated at all functions, meaning that in this case, the respective row of the design matrix would be coded as [0 0 0 1 0 0]. (C) After computing the betas and weighting the basis set by the estimated beta values, we obtain a staircase-like fit, clearly better than the linear predictor, but still poor. (D) Spline basis set. The range of the IV is covered by several spline functions that overlap with each other. Note that the example value of the IV (3.1°) produces non-zero values at several of the spline predictors (e.g. [0 0 0.5 0.8 0.15 0]). (E) After computing the betas and weighting the splines by the betas, we obtain a smooth fit.

Figure 5A shows a hypothetical nonlinear relationship between a predictor and the EEG. As an example, the predictor might be saccade amplitude and we see that the difference between a saccade of 1° and 2° has a much larger influence on the amplitude of the fixation-related lambda response than the difference between a 11° and 12° saccade. Obviously, a linear function would not fit this data well. An alternative way to model this relationship is to represent the continuous predictor, that is, the independent variable, by a set of basis functions (Figure 5B to 5E). One simple way to do this is to split up the range of the independent variable into multiple distinct bins (as used by Dandekar, Privitera, et al., 2012).

In the regression model, such a basis set is implemented by adding one additional column to the design matrix for each bin or basis function. For example, if we split the values of the saccade amplitude into six bins (Figure 5B), we would add five columns plus the intercept to the design matrix *X*. The independent variable now covers several columns and each column models a certain range of possible saccade amplitudes (e.g. 0-2°, 2-4°, 4-6°, …). When we solve the model, we estimate the beta weights for each basis function. As Figure 5D shows, this produces a better fit to the data as it captures the nonlinear relationship (Dandekar et al., 2012b). However, it also produces abrupt jumps between category borders, which can decrease statistical power and increase type-1 errors (Austin & Brunner, 2004). So instead, it is strongly recommended to keep the continuous predictors continuous (Altman & Royston, 2006; Bennette & Vickers, 2012; Collins, Ogundimu, Cook, Manach, & Altman, 2016; Royston, Altman, & Sauerbrei, 2006).

A better alternative, illustrated in panels D and E of Figure 5, is to use spline functions as a basis set. This is conceptually similar to the categorization approach but makes use of the assumption that the fit should be smooth. Instead of defining distinct bins, the range of possible values of the independent variable is covered by a number of spline functions that overlap with each other, as shown in Figure 5D (how exactly the spline set is constructed is outside the scope of this paper; the interested reader is referred to Wood, 2017).

If we now evaluate this set of splines at a given value of the independent variable (e.g. for a saccade of 3.1°), we obtain non-zero values not just for a single predictor, but for several neighboring spline functions as well (note that three functions are non-zero in Figure 5D). The different splines will have different strengths as they have different amounts of overlap with this value. When we solve the model, the spline functions are again weighted by their respective beta coefficients and summed up to obtain the modeled values. In contrast to the categorization approach, the result is a smooth, nonlinear fit (Figure 5E). In practice, however, we are not interested in the betas for each of the individual spline functions; instead, we want to evaluate the whole set of overlapping splines at certain values of the independent variable. For example, to visualize the effect of saccade amplitude on FRPs (cf. Figure 7E), we might want to evaluate the spline set, weighted by the previously estimated betas, at several saccade sizes of interest (e.g. 0.6°, 5°, 10°, and 15°).

When modeling nonlinear effects, one important parameter to set is the number of splines that cover the range of the predictor, because this determines how flexible (or “wiggly”) the modeled relationship can be. Using too few splines or too many splines increases the risk of underfitting or overfitting the data, respectively. Several automatic selection methods exist to set the number of splines, which are usually based on cross-validation or penalized fitting (Wood, 2017). Unfortunately, in case of the large deconvolution models, methods such as cross-validation are computationally very expensive. Also, while it is still possible to use crossvalidation to determine the best number of splines for a single – or just a few – spline predictors (e.g. with the help of function *uf_checkmodelfit.m* in the unfold toolbox), the problem quickly becomes intractable when the model contains many spline predictors and the optimal number of splines needs to be determined for each predictor. A pragmatic approach that we currently employ is to err on the side of caution (that is, underfitting) by using a relatively small number of splines to model nonlinear relationships (i.e., five splines in all models presented here). For more discussions on this issu, the reader is referred to Ehinger & Dimigen (2019).

### *Analyzing a free viewing dataset with the* unfold *toolbox*

In the following, we will briefly go through the practical steps to run and analyze a deconvolution model with spline predictors, using the *unfold* toolbox and the hypothetical car/face experiment depicted in Figure 2. Parts of the following description are adapted from our toolbox paper (Ehinger & Dimigen, 2019). For a detailed technical documentation of the toolbox and its features, we refer the reader to that paper.

To begin, we need a continuous EEG dataset in the EEGLAB format (Delorme & Makeig, 2004) that also contains event markers (triggers) for the experimental events of interest (such as stimulus onsets). For free viewing experiments, we need additional events that code the onsets of saccade and/or fixations as well as the respective properties of these eye movements (e.g. amplitude and direction of saccades, duration and location of fixations, the type of object fixated). With existing open source software, such eye movement events can be easily imported or detected in the synchronized eye tracking data. In most cases, the EEG data that we wish to analyze should have already been preprocessed (e.g. filtered) and corrected for ocular artifacts, for example with ICA procedures.

We then start the modeling process by writing down the model formula, which defines the design matrix *X* of the regression model. In the *unfold* toolbox, models are specified using the common Wilkinson notation (Wilkinson & Rogers, 1973) that is also used in other statistics software like *R*. Using this notation, we might define the following model for the hypothetical free viewing experiment depicted in Figure 2, during which participants looked at cars and faces:

~~~
                              FRP ~ 1 + cat(is_car) + sacc_amplitude
~~~

Here, the FRP is modeled by an intercept term (`1`) that describes the overall waveform, by a categorical variable (or factor) `is_car` that codes whether the currently looked at object is a car (1) or a face (0), and by a continuous linear predictor that codes the amplitude of the saccade that precedes fixation onset. It is also possible to define interactions (e.g., `is_car * sacc_amplitude`).

As explained earlier, it is unrealistic to always assume a linear influence of oculomotor behavior on the EEG. We can therefore relax this assumption and model saccade amplitude as a nonlinear predictor. With the following formula, the effect of saccade amplitude would be modeled by a basis set consisting of five splines ^1^:

~~~
                           FRP ~ 1 + cat(is_car) + spl(sacc_amplitude,5)
~~~

In the same model, we can simultaneously model brain responses evoked by other events, such as stimulus onsets or button presses. Each of these other event types can be modeled by their own formula. For example, in the car/face task, it would be important to also model the stimulus-ERP that is elicited by the onset of the car/face display on the screen. Otherwise this stimulus-long ERP will distort the baseline intervals of the following FRPs. This issue is crucial in experiments in which stimuli belonging to different conditions are fixated at slightly different average latencies after stimulus onset (Coco et al., 2020). For example, if the first fixation in a trial is aimed more often at a face than at a car, the face-FRP will be distorted differently by the overlapping stimulus-locked waveform that the car-ERP. Fortunately, the ERP evoked by the stimulus presentation can be simply accounted for by adding an additional intercept model for all stimulus events. In this way, it will be removed from the estimation of the FRPs. The complete model would then be:

~~~
                                         ERP ~ 1 {for stimulus onsets events}

                       FRP ~ 1 + cat(is_car) + spl(sacc_amplitude,5) {for fixation onset events}
~~~

Once the formulas are defined, the design matrix *X* is time-expanded to X_*dc*_ and now spans the duration of the entire EEG recording. Subsequently, the equation (EEG = X_dc_ * *b* + e) is solved for *b*, the betas. This is done for each channel separately. The resulting regression coefficients, or betas, correspond to the subject-level ERP waveforms in a traditional ERP analysis (Smith & Kutas, 2015a). For example, in the model above, for which we used the default treatment coding in the unfold toolbox, the intercept term of the FRP corresponds to the average FRP at the subject level elicited by a face fixation. The other betas, for `cat(is_car)`, will capture the partial effect of that particular predictor – here the effect of fixating a car rather than a face – and therefore corresponds to a difference wave in a traditional ERP analysis (here: car-FRP minus face-FRP). For data visualization or for second-level statistical analyses at the group level, these regression-based waveforms can therefore be treated just like any other subject-level ERP. In the following, we will apply this approach to several real datasets.

## EXPERIMENTAL METHODS

The empirical part of this paper will demonstrate the possibilities and advantages of (non)linear deconvolution modeling for analyzing free viewing experiments. For this, we will use co-registered data from three commonly studied paradigms: face recognition, scene viewing, and natural sentence reading. The following section briefly summarizes the experimental methods that are common to all three experiments.

### Participants

Participants in all three co-registration experiments were young adults, mostly psychology students, with normal or corrected-to-normal visual acuity (verified using Bach, 2006). Different participants took part in the three studies. We analyzed 10 participants for the face recognition experiment, 10 for the scene viewing experiment, and 42 for the sentence reading study. Experiments were conducted in compliance with the declaration of Helsinki (2008) and participants provided written informed consent before participation.

### Apparatus & Eye-Tracking

All datasets were recorded in an electromagnetically shielded laboratory at Humboldt-University using identical eye-tracking, EEG, and stimulation hardware. In all experiments, stimuli were presented at a viewing distance of 60 cm on a 22-inch CRT monitor (Iiyama Vision Master Pro 510, resolution 1024 × 768 pixels, vertical refresh 100 or 160 Hz depending on the experiment). Binocular eye movements were recorded at a rate of 500 Hz with a table-mounted eye tracker (IView-X Hi-Speed, SMI GmbH) that was frequently (re-)calibrated using a 9-point or 13-point grid and validated on a 4-point grid. Stimulus presentation and recordings were controlled by *Presentation* software (Neurobehavioral Systems, Albany, CA). Saccades and fixations were detected offline using the algorithm of Engbert & Kliegl (2003) as implemented in the EYE-EEG toolbox.

### Electrophysiological recordings

Electrophysiological signals were recorded from either 46 (*Face* and *Scene* experiment) or 64 (*Reading* experiment) Ag/AgCl electrodes. EEG electrodes were mounted in a textile cap at standard 10-10 system positions. Electro-oculogram (EOG) electrodes were positioned at the outer canthus and infraorbital ridge of each eye. Data were amplified with BrainAmp amplifiers (Brain Products GmbH) and digitized at 500 Hz, with impedances kept below 5 kΩ. Electrodes were initially referenced against an electrode on the left mastoid bone, but digitally re-referenced to an average reference. The *Face* and *Scene* data was acquired with a time constant of 10 s, whereas the *Reading* data was acquired as DC data. Offline, data of all experiments were high-pass filtered at a cutoff (−6 dB) of 0.1 Hz using EEGLAB’s finite response windowed sinc filter with default settings. Datasets were also low-pass filtered at 100 Hz (*Face* experiment) or 40 Hz (*Reading* and *Scene* experiment) using the same function.

EEG and eye-tracking were synchronized offline using EYE-EEG (version 0.81) based on shared trigger pulses sent from the presentation computer to the computers recording EEG and eye movements. The mean synchronization error was < 1 ms, computed based on the trigger alignment. Proper signal synchronization was additionally verified by computing the cross-correlating function between the horizontal gaze position signal and the horizontal bipolar electrooculogram which consistently peaked at or near lag zero (function *checksync.m* in EYE-EEG).

EEG data from the *Scene* experiment were corrected for ocular artifacts using Infomax ICA that was trained on band-pass filtered training data (Dimigen, 2020). Ocular components were then removed using the eye-tracker-guided method proposed by Plöchl et al., 2012 (variance ratio threshold: 1.1). Data for the reading experiment were artifact-corrected using Multiple-Source Eye Correction (Berg & Scherg, 1994) as implemented in BESA (Besa GmbH). Data of the *Face* experiment were not corrected for ocular artifacts, because there was a central fixation instruction during this experiment and the data therefore only contained comparatively small saccades. Instead, for this dataset, artifact-contaminated intervals were identified offline by moving a window with a length of 2000 ms in steps of 100 ms across the continuous recording. Whenever this window contained a peak-to-peak voltage difference > 150 μV at any channel, the corresponding EEG interval was removed from the deconvolution model. This is accomplished by setting all columns of the time-expanded design matrix (*X_dc_*) to zero for these “bad” time intervals; this way, they will be ignored in the regression model.

### Statistics

Statistical comparisons in all experiments were performed using the threshold-free cluster-enhancement method (Mensen & Khatami, 2013; S. M. Smith & Nichols, 2009), a data-driven permutation test that controls for multiple comparisons across time points and channels (using ≥ 1000 random permutations).

## EXPERIMENT 1: FACE PERCEPTION

In typical EEG experiments, participants are instructed to avoid eye movements. Yet, high-resolution eye-tracking reveals that even during attempted fixation, the eyes are not completely motionless, but frequently perform microsaccades, small involuntary movements with a typical amplitude of less than 1° (Rolfs, 2009). Depending on the task, these microsaccades are often found intermixed with small exploratory saccades aimed at informative regions of the stimulus, for example the eye region of a face. In the following, we will refer to both kinds of small eye movements simply as “miniature saccades” (Yuval-Greenberg et al., 2008).

Previous co-registration studies have shown that despite their small size, miniature saccades can generate sizable eye muscle (Craddock et al., 2016; Yuval-Greenberg & Deouell, 2009) and brain potentials (Dimigen et al., 2009; Gaarder et al., 1964; Meyberg et al., 2015) in EEG. Furthermore, because the amplitude and rate of miniature saccades often differs systematically between experimental conditions (Rolfs, 2009), these additional signals can seriously distort stimulus-locked analyses in the time and frequency domain (Dimigen et al., 2009; Yuval-Greenberg et al., 2008). In the following, we will demonstrate how deconvolution and spline regression can be used to elegantly control for the effects of miniature saccades and improve the data quality even in standard ERP experiments in which participants are told to maintain fixation.

### Participants

Twelve participants took part in the study. Here we analyze the data of 10 participants (19-35 years old, 8 female), because the data of two participants could not be synchronized across the whole recording. Single-subject data from one participant of this study was also shown in Ehinger & Dimigen (2019).

### Methods

During this experiment, previously described in the supplement of Dimigen et al. (2009), participants saw 480 pictures of human faces (7.5° × 8.5°) with a happy, neutral, or angry facial expression. The participants’ task was to classify the face’s emotional expression using three manual response buttons. At the start of each trial, a small (0.26°) central fixation cross appeared for 1000 ms. It was then replaced by the face for 1350 ms. Afterwards, the fixation cross re-appeared. Before the experiment, participants received written instruction to keep their fixation at the screen center for the duration of each trial.

We synchronized the EEG and eye-tracking data and then specified the following model: The formula for the stimulus onset events was `ERP ~ 1`, meaning that the stimulus-ERP was modeled by a constant intercept term (the factor “emotion” was ignored for the current analysis). Potentials from miniature saccades were modeled by the formula `Saccade-ERP ~ 1 + spl(sacc_amplitude,5)`, that is, by a constant intercept term and by a predictor coding saccade amplitude. Because the effect of saccade amplitude on the post-saccadic brain response is nonlinear, this effect was modeled here by five splines. Neural responses were estimated in the time window between −500 and 800 ms around each stimulus/saccade event and subsequently baseline corrected from −100 to 0 ms.

### Results & Discussion

Results are shown in Figure 6. Eye-tracking revealed that participants made at least one miniature saccade in the vast majority (99%) of trials. With a median amplitude of 1.5° (Figure 6B), most of these saccades were not genuine microsaccades, but rather small exploratory saccades aimed at the eyes or at the mouth region (Figure 6A), the parts of the face most informative for the emotion classification task.

**Figure 6.**
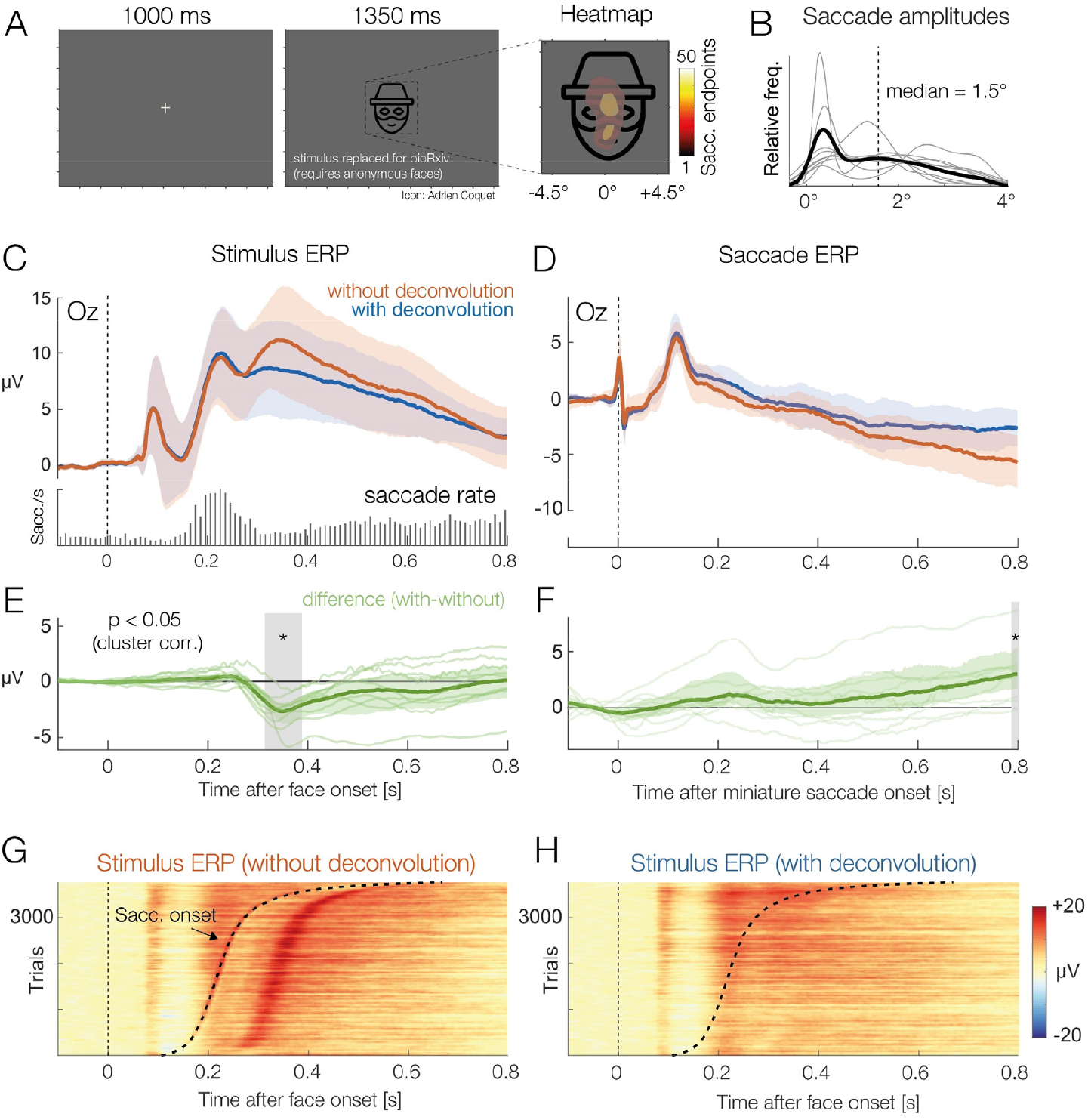
Grand-average results of the face classification experiment, with and without deconvolution. (A) On each trial, participants saw one face stimulus. Despite the instruction to avoid eye movements, participants made miniature saccades in almost every trial. Saccade endpoints are visualized as a heatmap, superimposed on the average face stimulus. (B) Miniature saccades had a median amplitude of 1.5°. Thin gray lines depict results for individual participants. (C) ERP at occipital electrode Oz, aligned to stimulus onset, without and with deconvolution (mean ±95% bootstrapped confidence intervals). The embedded histogram plots the saccade rate, that is, the occurrences of miniature saccades on the face. (D) Brain potentials time-locked to the onsets of miniature saccades detected during the trials. (E) Difference between the stimulus-ERP with and without deconvolution. The largest ERP difference can be seen about 100 ms after the rate of saccades reaches its maximum. (F) Same difference, but for the saccade-related ERP. (G) Color-coded single-trial EEG (*erpimage*) time-locked to face onset. When single trials are sorted by the latency of the first miniature saccade in the trial the overlapping saccade-related activity becomes evident. Note how the distortion of the stimulus ERP (in panel C and E) can be explained by the peak in saccade rate around 230 ms and the resulting overlap with the post-saccadic lambda response (shown in D). (H) Same data as in panel G, but after deconvolution. The *erpimage* also includes the residuals, that is, the EEG activity not explained by the model. Saccade-related activity was successfully removed.

The histogram in the lower part of Figure 6C shows that the rate of miniature saccades reached a maximum around 240 ms after stimulus onset. Each miniature saccade elicits its own visually-evoked lambda response (Dimigen et al., 2009) which peaks around 110 ms after saccade onset. Therefore, we would expect an impact of saccades on the stimulus-ERP beginning around 350 ms (240 + 110 ms) after stimulus onset. Indeed, if we compare the stimulus-ERP with and without deconvolution at occipital electrode Oz (panel C), we see a positive shift in the uncorrected signal that starts around 350 ms and continues until the end of the analysis time window.

Figure 6D shows the saccade-ERP, again with and without deconvolution. As expected, the saccadic response was also changed by the deconvolution, because we removed from it the overlapping influences of the stimulus-ERP as well as those other miniature saccades. Similar results have recently been reported by Kristensen et al. (2017, their Figure 6).

This simple example shows how linear deconvolution can disentangle eye movement-related potentials from stimulus-evoked activity to obtain an unbiased version of the stimulus-ERP not contaminated by saccade evoked activity. Deconvolution is especially important in experiments in which the rate, direction, or amplitude of saccades differs between conditions (Dimigen et al., 2009; Engbert & Kliegl, 2003; Meyberg, Sommer, & Dimigen, 2017; Yuval-Greenberg et al., 2008). However, even in cases where oculomotor behavior is the same in all conditions, the signal-to-noise ratio of the stimulus-locked ERP should improve after removing the brain-signal variance produced by miniature saccades (as already suggested in one of the earliest EEG/eye-tracking papers, Armington et al., 1967). Another major advantage compared to traditional averaging is that deconvolution also provides us with a clean, unbiased version of the (micro)saccade-related brain activity in the task. This is interesting, since potentials from small saccades have been shown to carry valuable information about the time course of attentional (Meyberg et al., 2015) and affective (Guérin-Dugué et al., 2018) processing in the task. With deconvolution, we can mine these “hidden” brain responses to learn more about the participant’s attentional or cognitive state.

## EXPERIMENT 2: SCENE VIEWING

Next, we will model fixation-related activity during natural scene viewing. As explained in the Introduction, the properties of eye movement-related brain potentials are not yet fully understood. What is clear, however, is that in addition to local stimulus features, properties of the incoming saccade strongly influence then neural response following fixation onset (Armington and Bloom, 1974; Thickbroom et al., 1991). This means that even a slight mismatch in oculomotor behavior between two conditions will produce spurious differences between the respective brain signals. Fortunately, deconvolution modeling can simultaneously control for overlapping potentials and the effects of oculomotor (e.g. saccade size) and visual (e.g. foveal image luminance) low-level variables.

In the following, we model FRPs from a visual search task on natural scenes. For the sake of clarity, in this example, we will only focus on the results of two oculomotor variables: saccade amplitude and saccade direction. These variables are highlighted here, because as we will show below, they both have clearly nonlinear effects on the FRP. Previous studies have already reported nonlinear influences of saccade amplitude on FRPs (Dandekar, Privitera, et al., 2012; Dimigen et al., 2009; Kaunitz et al., 2014; Ries et al., 2018b; Thickbroom et al., 1991; Yagi, 1979). For example, when executed on a high-contrast background, microsaccades of just 0.3° generate a lambda responses (P1) that is almost as large as that following a 4.5° saccade (cf. Figure 2 in Dimigen et al., 2009). Results like these suggest that it is insufficient to model saccade amplitude as a linear predictor. Effects of saccade direction on the post-saccadic neural response have only been recently reported (Meyberg et al., 2015, Cornelissen et al., 2019). As we show below, the direction of the preceding saccade has indeed a significant and nonlinear effect on the FRP during scene viewing.

### Participants

Ten young adults (age: 19-29 years, 4 female) with normal or corrected-to-normal visual acuity participated in the study.

### Methods

In the experiment, participants searched for a target stimulus hidden within images of natural scenes. Scenes consisted of grayscale versions of the first 33 images of the *Zurich Natural Image Database* (Einhäuser, Kruse, Hoffmann, & König, 2006), a collection of photographs taken in a forest (see Figure 7A for an example). Scenes spanned 28.8° × 21.6° of visual angle (800 × 600 pixels) and were centered on an otherwise empty black screen (resolution: 1024 × 768 pixel). One image was shown on each trial. The participant’s task was to find a small dark gray dot (0.4 cd/m^2^) that appeared at a random location within the scene at a random latency 8-16 seconds after scene onset. At first, the dot appeared with a diameter of just 0.07°, but then gradually increased in size over the course of several seconds. Once participants found the target, they pressed a button, which terminated the trial. A full manuscript on this dataset is currently in preparation.

**Figure 7.**
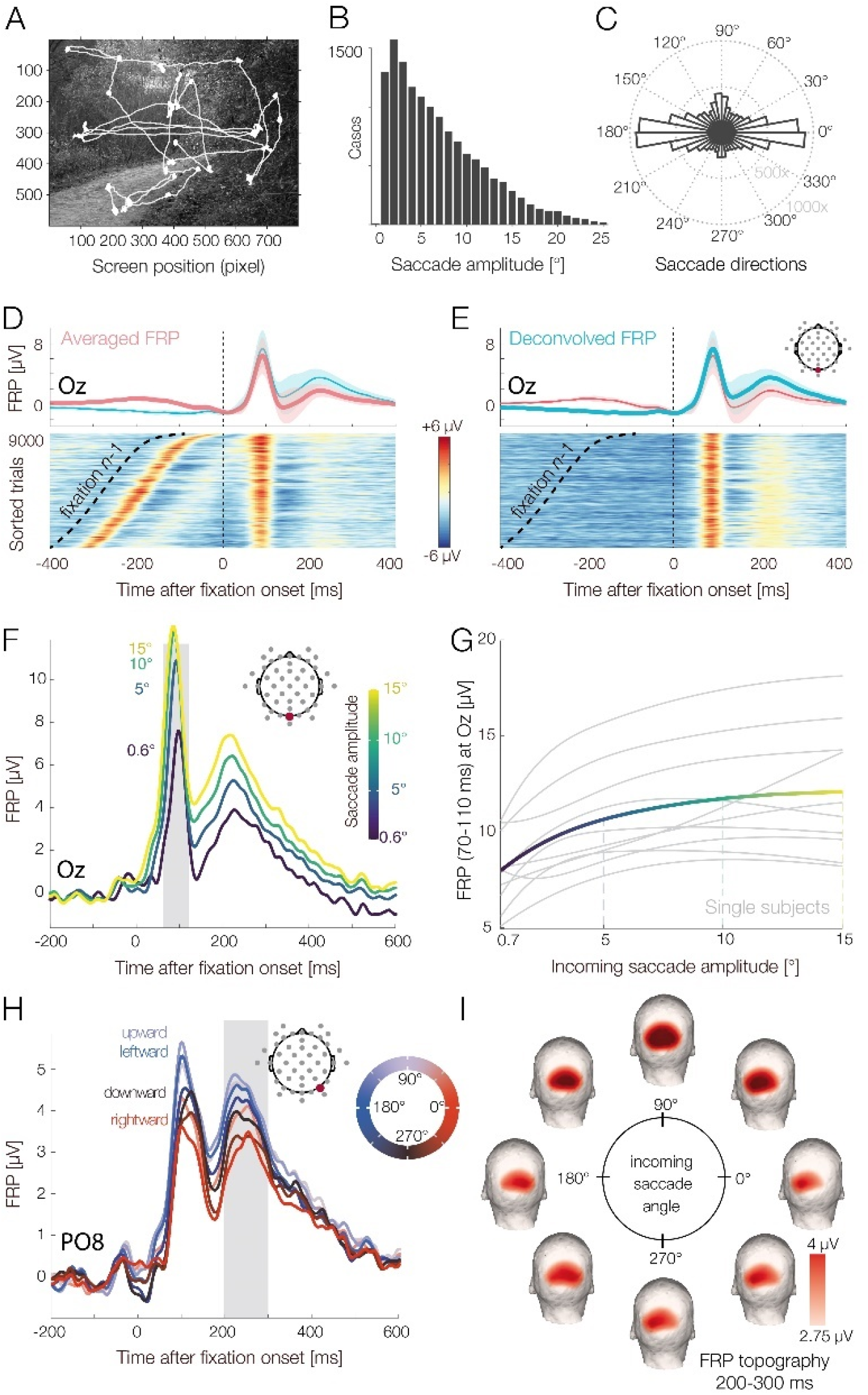
Visual search experiment with natural scenes. (A) Example scene with overlaid gaze path. (B) Distribution of saccade amplitudes and (C) saccade directions in the task. (D) FRP waveform over visual cortex without deconvolution (simple average, red line) and with deconvolution (blue line). Color-coded “single-trial” EEG of all participants (*erpimage*), sorted by the onset latency of the preceding (*n*-1) fixation on the scene. (E) Same FRP data as in (D). The *erpimage* shows the estimated deconvolved response plus the residuals, that is, the EEG activity not explained by the model. The overlapping activity from fixation *n*-1 was successfully removed. (F) Effect of saccade amplitude on the FRP, as revealed by the deconvolution model. Saccade amplitude modulates the entire post-saccadic waveform, with EEG amplitude changing as a nonlinear function of saccade size for the P1 (lambda response) and as a more linear function for the N1 component. (G) When saccade size is plotted against P1 amplitude (within the gray-shaded interval in panel F), the nonlinear, logarithmic relationship becomes obvious. The solid line depicts the mean over participants; gray lines depict single-subject estimates. (H) The FRP is also affected by the direction of the preceding saccade, with upwards saccade generating the strongest brain responses. This effect is again nonlinear and was therefore modeled by a set of circular spline functions coding the saccade angle. (I) Scalp topographies of the saccade direction effect at the latency of the second positive deflection (P2, 200-300ms, see gray shading in panel H). A topographic lateralization for leftward versus rightward saccades and a differentiation between upward versus downward saccades can be seen. Electrode PO7, on the left side of the head, showed a mirror-symmetric pattern.

For analysis, we specified the following model, which includes two types of events, the stimulus onset at the beginning of each trial and the onsets of fixations on the scene:

~~~
                                           ERP ~ 1

            FRP ~ 1 + spl(fixation_position_x,5) + spl(fixation_position_y,5) +
                  spl(sacc_amplitude,5) + circspl(sacc_angle,5,0,360)
~~~

For the stimulus-ERP we included only an intercept term, which captures the long-lasting ERP evoked by the presentation of the scene. Including it in the model ensures that FRPs are not distorted by the overlap with this stimulus-ERP. For the fixation onset events we modeled the horizontal and vertical fixation position on the screen as well as the incoming saccade amplitude using spline predictors. In addition, we modeled the direction of the incoming saccade. Because the angle of a saccade is a circular predictor, ranging from 0 to 360 degrees of angle, it was modeled by a set of five circular splines (Ehinger & Dimigen, 2019). Responses were modeled in a time window from −400 to 800 ms around each event.

To sum up the model, we modeled the ERP elicited by the scene onset and the FRPs elicited by each fixation on the scene and allowed for several nonlinear effects of saccade properties on the FRP. In the following, we will focus on the results for two of these predictors: saccade amplitude and saccade angle.

### Results & Discussion

Panels B and C of Figure 7 summarize the eye movement behavior in the task. Saccades had a median amplitude of 4.9° and fixations lasted on average 264 ms. Electrophysiological results are summarized in Figure 7D to 7I.

Panels D and E of Figure 7 compare the raw FRP (obtained with simple averaging, red line) to the regression-FRP obtained with the deconvolution model (blue line). The *erpimages* at the bottom of each panels show the corresponding EEG data at the single-trial level. To illustrate the impact of overlapping activity, single-trial epochs were sorted by the onset latency of the preceding fixation (*n*-1) on the scene. In the raw data (Figure 7D), it is obvious that the neural activity from fixation *n*-1 distorts the waveshape of the current fixation *n*, especially during the baseline interval. In the modeled data, which also includes the residual activity not accounted for by the model (Figure 7E), activity aligned to fixation *n*-1 is no longer visible. Importantly, the lack of residual activity time-locked to fixation *n*-1 suggests that the neural activity during scene viewing was successfully modeled and corrected for overlapping activity.

Panels F and G of Figure 7 show the partial effects of saccade amplitude and saccade direction taken from the deconvolution model. The isolated effect of saccade amplitude (Figure 7F and 6G) reveals a long-lasting impact of saccade size on the FRP waveform: At electrode Oz, located over primary visual cortex, saccade amplitude influenced all time points of the FRP up to 600 ms after fixation onset. Results also confirm that this effect is indeed highly nonlinear: the increase in P1 amplitude with saccade size was steep for smaller saccades (< 6°) but then slowly leveled off for larger saccades. Such nonlinearities were observed for all ten participants (Figure 7F). It is obvious that a nonlinear model is more appropriate for this data than a linear one.

Interestingly, the angle of the incoming saccade also modulated the FRP in a highly nonlinear manner. In Figure 7G, this is shown for lateralized posterior electrode PO8, located over the right hemisphere. The corresponding scalp topographies for saccades of different directions are shown in Figure 7H, in the time window 200 to 300 ms after fixation onset. It can be seen how saccade direction changes the FRP’s scalp distribution, with rightward-going saccades generating higher amplitudes over the left hemisphere and vice versa (see Cornelissen et al., 2019; and Meyberg et al., 2015 for similar findings). Note that this effect is not due to corneoretinal artifacts, which were successfully suppressed with ICA. This effect of saccade direction is also not explained by different fixation locations on the screen following saccade offset (Dimigen et al., 2013), since horizontal and vertical fixation position were also included as predictors in the model (results not shown here).

### Discussion

In this example, we simultaneously modeled the effects of some oculomotor covariates on FRPs. During scene viewing, these low-level covariates are often intercorrelated with each other (Nuthmann, 2017) and correlated with the high-level “cognitive” factors of interest (e.g. whether the fixated item is the search target). Furthermore, as we show here, they influence long intervals of the FRP waveform in a nonlinear way. A strictly linear model (Coco et al., 2020; e.g. linear saccade amplitude in Cornelissen et al., 2019; Kristensen, Guerin-Dugué, et al., 2017; Weiss et al., 2016) is therefore not ideal to capture these complex relationships.

In addition to the covariates discussed here, one could easily enter more low-level predictors into the model, such as the local luminance and power spectrum of the currently foveated image region. Finally, to study cognitive effects on the fixation-related P300 component in this task (Dandekar, Ding, Privitera, Carney, & Klein, 2012; Dias et al., 2013; Kamienkowski et al., 2012; Kaunitz et al., 2014), one could add a categorical predictor (0 or 1) coding whether the fixated screen region contains the search target participants were looking for. The next example will illustrate how we can reliably study the time course of such psychologically interesting effects during active vision.

## EXPERIMENT 3: NATURAL READING

In free viewing experiments, the psychological manipulation of interest is typically linked to a change in fixation duration, which will distort the FRPs. A classic task to illustrate this problem and its solution via deconvolution modeling is reading. In ERP research on reading, sentences are traditionally presented word-by-word at the center of the screen. While this serial visual presentation procedure controls for overlapping potentials, it differs in important ways from natural reading (Kliegl, Dambacher, Dimigen, & Sommer, 2014; Sereno & Rayner, 2003). One key property of visual word recognition that is neglected by serial presentation procedures is that the upcoming word in a sentence is usually previewed in parafoveal vision (eccentricity 2-5°) before the reader looks at it (Schotter, Angele, & Rayner, 2012). The parafoveal preprocessing then facilitates the recognition of the word once the word is fixated. This facilitation is evident in the classic *preview benefit* (Rayner, 1975) in behavior: words that were visible during preceding fixations receive 20-40 ms shorter fixations (Vasilev & Angele, 2017) than words that were gaze-contingently masked with a different word or a meaningless string of letters before being fixated.

Combined eye-tracking/EEG studies have recently established tentative neural correlates of this preview benefit in FRPs: An early effect, corresponding to a reduction of the late parts of the occipitotemporal N1 component between about 180-280 ms after fixation onset (“preview positivity”) that is sometimes followed by a later effect at around 400 ms that may reflect a reduction of the N400 component by preview (Degno et al., 2019; Dimigen, Kliegl, & Sommer, 2012; Kornrumpf, Niefind, Sommer, & Dimigen, 2016; Li, Niefind, Wang, Sommer, & Dimigen, 2015; Niefind & Dimigen, 2016). However, an inherent problem with all previously published studies is that the difference in fixation times measured on the target word also changes the overlap with the following fixations. This raises the question to which degree the reported neural preview effects are real or just a trivial consequence of the different overlap situation in the conditions with and without an informative preview. Below we will demonstrate how deconvolution modeling can answer this question by separating genuine preview effects from spurious overlap effects.

### Participants

Participants were native German speakers with normal or corrected-to-normal visual acuity (mean age: 25.7 years, range 18-45 years, 27 female). Here we present results from the first 42 participants recorded in this study. A manuscript describing the full dataset is currently in preparation^2^.

### Methods

Participants read 144 pairs of German sentences belonging to the *Potsdam Sentence Corpus 3*, a set of materials previously used in psycholinguistic ERP research and described in detail in Dambacher et al. (2012). On each trial, two sentences were successively presented as single lines of text on the monitor (Figure 8A). Participants read the sentences at their own pace and answered occasional multiple-choice comprehension questions presented randomly after one third of the trials. Sentences were displayed in a black font (Courier, 0.45° per character) on a white background. The second sentence contained a target word (e.g. “weapon”) for which the parafoveal preview was manipulated using Rayner’s (1975) boundary paradigm (Figure 8A): Before fixating the target word, that is, during the preceding fixations, readers either saw a correct preview for the target word (e.g. “weapon”) or a non-informative preview that consisted of a meaningless but visually similar letter string of the same length (e.g. “`vcrqcr`”). During the saccade to the target, while visual sensitivity is reduced (Matin, 1974), the preview mask was always gaze-contingently exchanged with the correct target word (e.g. “`weapon`”). This display change was executed with a mean latency of < 10 ms and typically not noticed by the participants, as validated with a questionnaire after the experiment.

**Figure 8.**
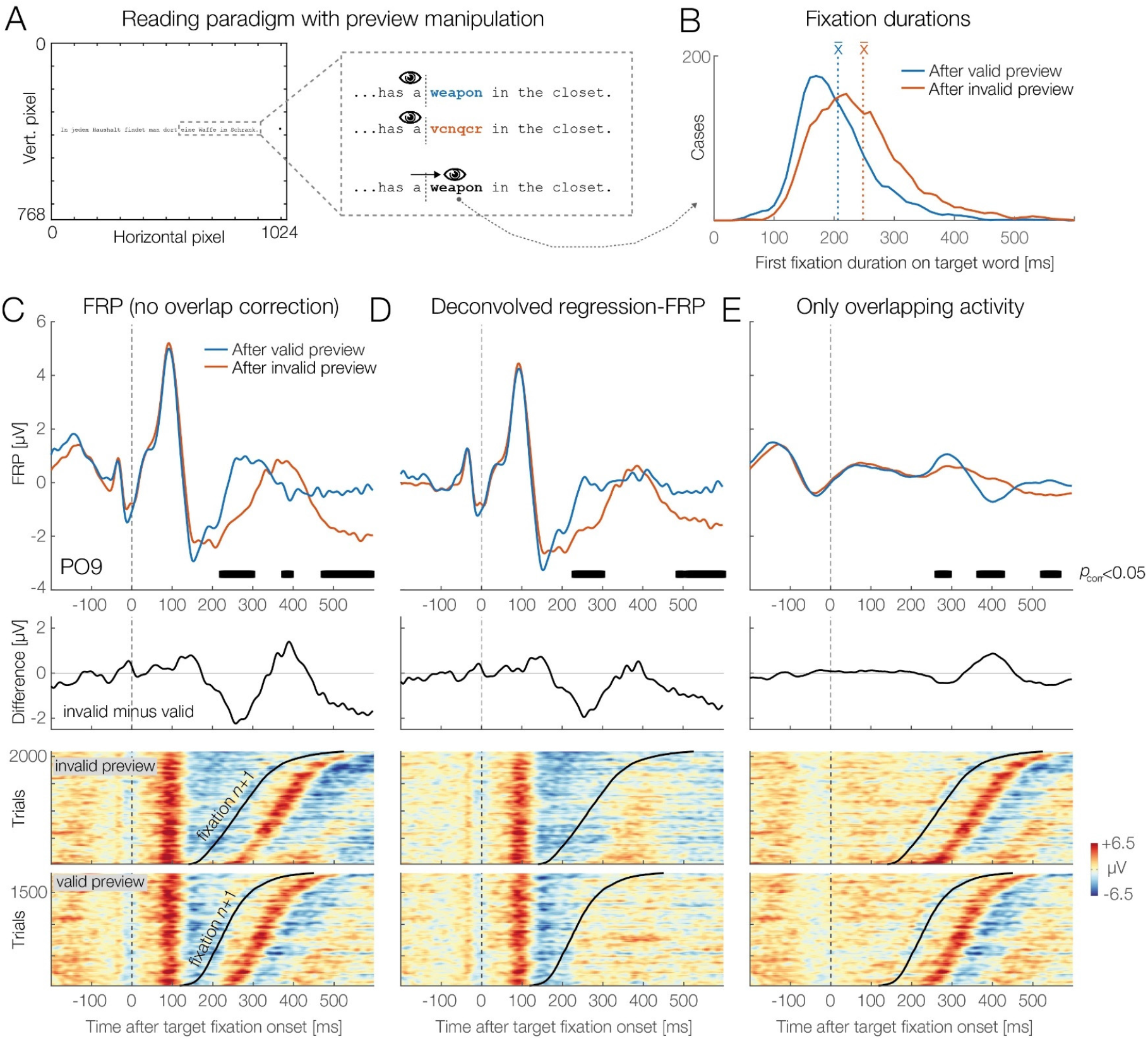
Grand-average results from the sentence reading experiment (*N*=42). (A) Participants read sentences from left to right. For one target word, the parafoveal preview was manipulated with the boundary technique. (B) Preview benefit in behavior. Fixations on target (e.g., “weapon”) were 41 ms shorter when the parafoveal preview shown during the preceding fixation was valid (correct word, highlighted here in blue) rather than invalid (random letter preview, red). (C) Grand-average FRP at occipitotemporal electrode PO9, aligned to the first fixation on the target. Black bars indicate the extent of the clusters that contributed to the overall significant effect of preview on the FRP. Lower panel: Color-coded single-trial epochs, sorted by the fixation duration on the target word, reveal that the FRP waveform is a mixture of potentials evoked by the current and the next fixation. (D) Same data, but corrected for overlapping potentials. We can see that the middle cluster (from about 382-400 ms) has disappeared, because it was only caused by the different overlap situation in the valid/invalid condition. In contrast, the clusters around 228-306 ms (“preview positivity”, Dimigen et al. 2012) and 480-600 ms remain visible because they are not a trivial result of overlap. (E) The isolated overlapping activity from neighboring fixations differed significantly (*p* < 0.05) between conditions. In addition to modulating the effect sizes of the early and late preview effect, overlap also produced the spurious middle-latency cluster (around 390 ms).

To analyze the data, we first marked all trials that contained eye blinks, a loss of eyetracking data, a late display change (executed > 10 ms after saccade offset), a skipping of the target word, or excessive non-ocular EEG artifacts. Remaining non-ocular artifacts were detected by shifting a 1000 ms long moving window across the continuous EEG and by marking all intervals in which the window contained peak-to-peak voltage differences > 120 μV in any channel. In the deconvolution framework, these “bad” intervals can then be easily excluded by setting all columns of the design matrix to zero during these intervals (Smith & Kutas, 2015b). The mean number of remaining target fixations per participant was 51.0 (range: 36-65) for the invalid and 44.1 (range: 29-59) for the valid preview condition.

In the second step, we modeled both the ERP elicited by the sentence onset (with its intercept) as well as the FRP evoked by each reading fixation. The model specified was

~~~
                                              ERP ~ 1

                        FRP ~ 1 + cat(is_targetword) * cat(is_previewed)
                                     + spl(sac_amplitude, 5)
~~~

where `is_targetword` and `is_previewed` are both binary categorical predictors coding whether or not a fixation was on the manipulated target word and whether or not that target word had been visible during the preceding fixations, respectively. Saccade amplitude was again modeled by a spline predictor. We estimated the responses from −300 to +800 ms around each event and baseline-corrected the resulting waveforms by subtracting the interval from −100 to 0 ms. For group statistics, the interval between −300 and +600 ms after fixation onset at occipito-temporal channel PO9 was submitted to the TFCE permutation test.

### Results & Discussion

Table 1 reports the fixation durations in the target region of the sentence. The average duration of all fixations during sentence reading (including refixations) was only 207 ms, meaning that FRPs were strongly overlapped by those from preceding and subsequent fixations. Figure 8B visualizes the distribution of first-fixation durations on the target word (e.g. “weapon”) as a function of whether the preview for this word was valid (blue line) or invalid (red line). Note that the two conditions only differ in terms of what the participant saw as a preview during the *preceding* (pre-target) fixation; the foveated word is the same in both conditions. As expected, fixation durations on the pre-target word were not significantly affected by the preview manipulation. However, the subsequent first-fixation on the target word itself was on average 41 ms shorter in the condition in which a valid rather than an invalid preview was provided. This preview benefit was significant, *t*(41) = 10.82, *p* < 0.0001. In gaze duration, which is the summed duration of all first-pass fixations on the target word, the effect was also significant, with a difference of 51 ms, *t*(41) = 11.83, *p* < 0.0001. Results therefore replicate the classic preview benefit in behavior (Rayner, 1975).

**Table 1.** Fixation times (in ms) in the reading experiment

|  | First-fixation duration |  | Gaze duration |  |
| --- | --- | --- | --- | --- |
|  | Invalid preview | Valid preview | Invalid preview | Valid preview |
| Pre-target word | 216 (32) | 211 (28) | 238 (45) | 230 (44) |
| Target word | 248 (36) | 207 (24) | 275 (44) | 224 (30) |

Note: values in parentheses give one SD.

Figure 8C presents the corresponding FRP waveforms. The plotted channel is PO9, a left-hemispheric occipitotemporal electrode where the strongest preview effects have been observed previously (Dimigen et al., 2012; Kornrumpf et al., 2016). Time zero marks the fixation onset on the target word. Note that at this time, readers are always looking at the correct target word; conditions only differ in terms of what was parafoveally visible during the preceding fixation. As Figure 8C shows, permutation testing revealed a significant effect (*p* < 0.05, under control of multiple comparisons) of preview condition on the FRP after conventional averaging (without deconvolution). Black bars in panels 7C, 7D, and 7E highlight the duration of the underlying temporal clusters at electrode PO9. Please note that because these clusters are computed during the first stage of the permutation test (Sassenhagen & Draschkow, 2019) they are themselves not stringently controlled for multiple comparisons (unlike the overall test result). However, their temporal extent provides some indication of the intervals that likely contributed to the overall significant effect of preview. With conventional averaging, clusters extended across three intervals after fixation onset: early: 228-306 ms, middle: 382-400 ms, and late: 480-600 ms (black bars in Figure 8C). However, if we look at the underlying single-trial EEG activity sorted by the fixation duration on the target word (lower panel of Figure 8C), it becomes obvious that a relevant part of the brain potentials after 200 ms is not generated by the fixation on the target word but by the next fixation (*n+1*) within the sentence. Since this next fixation begins on average 41 ms later in the invalid preview condition (see Table 1), this creates a spurious condition difference in the FRP.

Figure 8D shows the same data, corrected for overlapping potentials. In the sorted singletrial data (lower panel), the activity related to the target fixation is preserved, whereas that from the next fixation *n*+1 is removed. Crucially, the overall significant effect of preview (*p* < 0.05) is preserved. However, instead of three, we now only observe two clusters, one extending from 232 to 300 ms (reflecting the early “preview positivity”) and another late one, beginning at 490 ms and lasting until the end of the epoch at 600 ms (possibly reflecting a preview effect on the N400). In contrast, the middle cluster around 390 ms has disappeared, because this difference was only caused by overlapping activity. This confounding effect of overlapping potentials is confirmed in Figure 8E, which shows just the activity produced by the neighboring non-target fixations. The permutation test confirmed that this overlapping activity alone produced a significant difference between conditions (*p* < 0.05) in three intervals (Figure 8E, black bars). In addition to modulating the strength of the genuine early and late preview effects, overlapping potentials produced the entirely spurious mid-latency effect at around 390 ms.

### Discussion

Existing studies on neural preview effects did not control for the signal distortions produced by the difference in fixation time. Our analysis confirms for the first time that the previously reported neural preview effects are not trivial artifacts of overlapping activity, but genuine consequences of parafoveal processing. This insight is important because during natural vision, most visual objects are already partially processed in the parafoveal or peripheral visual field before they enter the fovea. In other words, while ERP researcher have traditionally presented isolated objects during steady fixation, visual objects are typically primed and/or predicted during natural vision based on a coarse extrafoveal preview of the object. Indeed, a similar preview effect, as shown here for words, was recently reported for the N170 component of previewed human faces (Buonocore, Dimigen, & Melcher, 2020; de Lissa et al., 2019; Huber-Huber et al., 2018). This indicates that the attenuation of the late N1 and N170 components by preview may be a characteristic feature of visual object recognition under real-world conditions.

In summary, the application to reading demonstrates how deconvolution can disentangle cognitive effects from spurious overlap effects, allowing us to resolve the precise time course of mental processes within a single fixation.

## GENERAL DISCUSSION

Fixation-related potentials are a promising and data-rich tool to study the dynamics of visual cognition under ecologically valid conditions. In this article we have outlined an integrated analysis framework for combined eye-tracking/EEG experiments that integrates deconvolution modeling with nonlinear spline regression. Applications of this approach to three commonly-studied paradigms suggest that it can control for the confounding effects of overlap and low-level covariates on the FRP and provide new insights into cognitive processes during active vision.

In the face recognition study, our analysis confirmed that overlapping muscle and brain potentials from small involuntary saccades during the task are problematic and present in the majority of trials even in “traditional” EEG experiments. However, with deconvolution modeling, these saccade-related potentials could be effectively removed to obtain clean stimulus-locked ERPs. This also opens the interesting possibility to use the now isolated eye movement-related brain potentials as an additional source of information about the participant’s attentional, cognitive, or affective processing in the task (Meyberg et al., 2015; Guérin-Dugué et al., 2018). The scene viewing example was included to demonstrate that at least some of the (numerous) low-level influences on the FRP during free viewing are highly nonlinear. For example, our analysis revealed a nonlinear effect of the angle of the incoming saccade on the FRP. Finally, the application to natural reading showed how spurious effects due to different fixation durations can be removed. This allowed us for the first time to describe the time course of neural preview benefits in an unbiased manner. In contrast, the simple averaging approach used in previous studies (Degno et al., 2019; Dimigen et al., 2012; Kornrumpf et al., 2016) will have necessarily produced incorrect conclusions about the exact time course of this effect.

In the following, we will further discuss the underlying assumptions, possibilities, and existing limitations of using (non)linear deconvolution models for combined eye-tracking/EEG research and outline some interesting future perspectives.

### Assumptions of deconvolution models

Traditional ERP averaging is based on the assumption that the underlying event-related response is invariant across all epochs of a given condition and that the average is therefore a sensible description of the individual trials (Otten & Rugg, 2005). This assumption is likely incorrect, since cortical information processing likely varies between trials, as also indicated by trial-to-trial differences in reaction times. Deconvolution models are based on the same assumption, namely that the fixation-related response is the same, regardless of the amount of overlap. In other words, we rely on the somewhat unrealistic assumption that the neural response does not differ between short and long fixations. The assumption also concerns sequences of fixations. From ERP research, we know that the processing of one stimulus can change the processing of the next one due to adaptation, habituation, or priming (e.g. Schweinberger & Neumann, 2016). Again, it would be surprising if these factors do not also modulate the FRP while a stimulus is rapidly scanned with several saccades. If such sequential effects occur often enough in an experiment, they can be explicitly modeled within the deconvolution framework. For example, in a scene viewing study, on could add an additional predictor that codes whether a fixation happened early or late after scene onset (Fischer, Graupner, Velichkovsky, & Pannasch, 2013) or whether it was the first fixation or a refixation on a particular image region (Van Humbeeck et al., 2018).

### Baseline correction and placement

In ERP research, baseline correction is performed to accommodate for slow drifts in the signal, for example due to changes in skin potential (Luck, 2014). The baseline interval is therefore typically placed in a “neutral” interval immediately before stimulus onset. In experiments with multiple saccades, it is more difficult to find an appropriate neutral baseline (Dimigen et al., 2011; Nikolaev et al., 2016).

The first reason is of methodological nature and directly linked to the problems of overlap and covariates: The baseline for the FRP is often biased because of differences in the duration of the preceding fixation, differences in the size of the preceding saccade, or a different overlap with the stimulus-onset ERP. Several workarounds have been proposed to deal with this problem, such as placing the baseline before trial onset (Dimigen et al., 2011; Nikolaev et al., 2016) before an earlier fixation (Coco et al., 2018; Degno et al., 2018, Huber-Huber et al., 2018), or in the first few milliseconds after fixation onset (de Lissa et al., 2019; e.g. Simola, Fevre, Torniainen, & Baccino, 2014, see Nikolaev et al., 2016 for an illustration of different baseline placement options). With deconvolution, this is no longer necessary, because we can effectively remove the overlapping activity and covariate effects from the baseline interval. In our experience, the baselines of the deconvolved FRPs are essentially flat (as also visible in Figures 7E and 8D above). This means that a conventional baseline correction can be applied to the deconvolved FRPs.

However, a second reason why the baseline should still be chosen carefully are effects of preprocessing. Because viewers obtain some information about soon-to-be fixated items in parafoveal and peripheral vision (Baccino & Manunta, 2005; Buonocore et al., 2020; Coco et al., 2018; de Lissa et al., 2019), EEG effects may in some cases already begin before an object is foveated (e.g. Baccino & Manunta, 2005; Luo et al., 2009). In paradigms where such parafoveal-on-foveal effects are likely to occur, it may still be sensible to place the baseline interval further away from fixation onset, even after overlapping potentials are removed. An even better option would be to capture these parafoveal-on-foveal effects in the model itself by adding the pre-target fixations as a separate type of event to the model. For example, in the reading experiment reported above, we could have coded the status of each reading fixation (in the predictor `is_targetword`) not just as “non-target” fixation (0) or “target” fixation (1), but could have added a third category for “pre-target” fixations (those on the word before the target word). In this way, any potential parafoveal-on-foveal effects produced by seeing the parafoveal non-word mask during the pre-target fixation can be disentangled from the neural preview benefits after fixating the target.

### Time-frequency analysis

While most EEG recordings during free viewing have been analyzed in the time domain, it is also possible to study eye movement-related changes in oscillatory power and phase (Bodis-Wollner et al., 2002; Gaarder, Koresko, & Kropfl, 1966; Hutzler, Vignali, Hawelka, Himmelstoss, & Richlan, 2016; Kaiser, Brunner, Leeb, Neuper, & Pfurtscheller, 2009; Kornrumpf, Dimigen, & Sommer, 2017; Metzner, von der Malsburg, Vasishth, & Rösler, 2015; Nikolaev et al., 2016; Ossandon et al., 2010). Event-related responses in the frequency domain – such as induced changes in power – can last for several seconds and are likely biased by overlapping activity in much the same way as FRPs (Litvak et al., 2013; Ossandon et al., 2019). To address this problem, Ossandón and colleagues (Ossandón, König, & Heed, 2019) recently used the Hilbert transformation to obtain instantaneous power of the EEG in the alpha band. They then deconvolved the bandpass-filtered and rectified signal, showing that deconvolution can also be applied to EEG oscillations. Deconvolution is also an interesting option to correct for the spectral artifacts that are produced by involuntary microsaccades in time-frequency analyses (Yuval-Greenberg et al., 2008). Specifically, the results of the face recognition experiment (Figure 6D) suggest that deconvolution is able to isolate the saccadic spike potential, the eye muscle artifact that is known to produce strong distortions in the gamma band (> 30 Hz) of the EEG. Cognitive influences on stimulus-induced gamma oscillations can therefore likely be disentangled from microsaccade artifacts if the continuous gamma band power (rather than the raw EEG) is entered into the model.

An unresolved question concerns the most suitable measure of spectral power to put into the deconvolution model. Litvak and colleagues (2013) conducted simulations on this issue where they compared the model fits (R^2^) for different measures of spectral power (raw power, log power, square root of power) and obtained the best results for the square root of power. Further simulations are needed to see which of these transformations is most suitable or whether the differences are negligible in practice.

### Improving the understanding of fixation-related brain activity

There are many ways to further improve the estimation of FRPs during free viewing. For example, the lambda response, the predominantly visually-evoked P1 response following fixation onset, is not fully understood. Existing evidence suggests that it is itself a compound response, consisting of at least two separate subcomponents: a visual “off”-response produced by the beginning of the saccade and a visual “on”-response following the inflow of new visual information at saccade offset (Kazai & Yagi, 2003; Kurtzberg & Vaughan, 1982; Thickbroom et al., 1991). Potentially, deconvolution could be used to separate the saccade onset and saccade offset-related contributions to the lambda response. Another promising application is to isolate the possible neural correlates of peri-saccadic retinal stimulation, for example due to gaze-contingent display changes (Chase & Kalil, 1972; Kleiser, Skrandies, & Anagnostou, 2000; Skrandies & Laschke, 1997).

Another interesting feature of linear deconvolution is that it is possible to add temporally continuous signals – rather than only discrete event onsets – as predictors to the time-expanded design matrix (Gonçalves, Whelan, Foxe, & Lalor, 2014; Lalor, Pearlmutter, Reilly, McDarby, & Foxe, 2006). In the literature, this is typically referred to as the temporal response function approach (Crosse, Di Liberto, Bednar, & Lalor, 2016). This approach can be combined with the deconvolution approach summarized in the present paper (which uses discrete event onsets as regressors) and is also implemented in the *unfold* toolbox. For example, to partially correct for corneoretinal artifacts, one could add a column to the time-expanded design matrix that contains the continuous gaze position signal of the eye-tracker, which will then be regressed out from the EEG (as suggested by Dandekar et al., 2012b). Yet another possibility is to add the pupil diameter as a time-continuous predictor. The idea would be that the momentary level of arousal and mental load – as indexed by pupil size – will correlate with the amplitude of the neural response. Other continuous signals that could be added include the luminance profile of a continuously changing stimulus (e.g. of a video watched by the participant) or the sound envelope of concurrent auditory stimuli (e.g. the sound channel of the movie). Finally, signals from accelerometers and other motion sensors could help to account for the head and body movements that characterize visual exploration behavior outside of the laboratory (Ehinger et al., 2014; Gramann, Jung, Ferris, Lin, & Makeig, 2014).

### Integrating deconvolution with linear mixed models

In the present work, we used the two-stage statistical approach that is also commonly used with mass univariate models (Pernet et al., 2011). Here, the regression-ERPs (betas) are first computed individually for each participant and then entered into a second-level group analysis (e.g. an ANOVA or a permutation test). Compared to this hierarchical approach, linear mixed-effects models (e.g. Gelman and Hill, 2007) provide a number of advantages (Baayen et al., 2008; Kliegl et al., 2011), such as the option to include crossed random effects for subjects and items. Mixed models are often used to analyze fixation durations (e.g. Kliegl, 2007; Nuthmann, 2017; Ehinger et al., 2018) and more recently also FRPs (Dimigen et al., 2011; Degno et al., 2018). In the long term, it will be promising to integrate deconvolution with mixed-effects modeling (Ehinger, 2019), but this will require large computational resources (because the EEG data of all participants has to be fitted simultaneously) as well as new algorithms for estimating sparse mixed-effects models (e.g. Wood, Li, Shaddick, & Augustin, 2017).

### Towards a full analysis pipeline for free viewing EEG

In Figure 1, we had summarized four challenges that have complicated eye-tracking/EEG research in the past. We believe that there are now adequate solutions to all four problems. The *unfold* toolbox used for the current analyses is fully compatible with the existing EYE-EEG toolbox. In a first step, EYE-EEG can be used to synchronize the recordings, to add saccade and fixation events to the data, and to suppress eye movement artifacts with specialized ICA procedures (Dimigen, 2020, Plöchl et al., 2012). Afterwards, the artifact-corrected EEG can be read into the *unfold* toolbox to model the fixation-related responses. Taken together, the two toolboxes provide a full open source pipeline to address the four problems of co-registration.

### Conclusions

In this paper we have presented a framework for analyzing eye movement-related potentials and exemplified its advantages for three common paradigms. By controlling for overlapping potentials and low-level influences, the (non)linear deconvolution framework allows us to study new exciting phenomena that were previously difficult or impossible to investigate. In combination with existing approaches for data integration and artifact correction, this opens new possibilities to investigate natural viewing behavior with EEG without compromising data quality.

## Footnotes

1 The number of splines that cover the range of the predictor determines how flexible the fit is. A larger number of splines allows us to model more complex relationships but also increases the risk of overfitting the data. See section “Modeling non-linear effects” above and Ehinger & Dimigen (2019) for discussions.

2 The target word in the second sentence was also manipulated in terms of its contextual predictability and lexical frequency (Dambacher et al., 2012). Here we focus only on the factor preview.

